# Ribosomal RNA 2’-O-methylation dynamics impact cell fate decisions

**DOI:** 10.1101/2022.09.24.509301

**Authors:** SJ Häfner, MD Jansson, K Altinel, ML Kraushar, KL Andersen, Z Caldwell Abay-Nørgaard, M Fontenas, DM Sørensen, DM Gay, FS Arendrup, D Tehler, N Krogh, H Nielsen, A Kirkeby, AH Lund

## Abstract

Translational regulation impacts both pluripotency maintenance and cell differentiation. To what degree the ribosome itself exerts control over this process remains unanswered. Accumulating evidence has demonstrated heterogeneity in ribosome composition in various organisms. 2’-O-methylation of rRNA represents an important source of heterogeneity, where site-specific alteration of methylation levels can modulate translation. Here we explore changes in rRNA 2’-O-methylation during mouse brain development and during tri-lineage differentiation of human embryonic stem cells. We find distinct alterations between brain regions, as well as clear dynamics during cortex development and germ layer differentiation. We identify a methylation site which impacts neuronal differentiation. Modulation of its methylation levels affects ribosome association of the Fragile X Mental Retardation Protein and translation of WNT pathway-related mRNAs. Together, the data reveals ribosome heterogeneity through rRNA 2’-O-methylation during early development and differentiation and suggests a direct role for ribosomes in regulating translation during cell fate acquisition.

## Introduction

Embryonic development is known to require specific and accurately dosed protein subsets with utmost spatiotemporal precision, often paralleled by profound changes in cell proliferation and overall protein synthesis rates (Khajuria et al., 2018; Kraushar et al., 2015; Magee and Signer, 2021; R. Wang and Amoyel, 2022). In particular, the formation of the mammalian nervous system requires an exceptionally fine-tuned protein homeostasis in order to generate, organize, and connect hundreds of neural subtypes (Baser et al., 2019; Blair et al., 2017; Kapur et al., 2017; Kraushar et al., 2020), and any failure of the translation machinery derails normal brain development and function (Kapur et al., 2017; Laguesse et al., 2015; Lauria et al., 2020; Yamada et al., 2019). Hence, multiple layers of regulation converge during development to impact gene expression output. Whereas transcriptional, posttranscriptional, and posttranslational regulation have been studied in many developmental model systems (Mohammed et al., 2017; Y.-C. Wang et al., 2014; B. S. Zhao et al., 2017), the role of translation, and specifically the intrinsic regulatory potential of modifications to the ribosome itself, remains understudied (Baser et al., 2019; Lins et al., 2022).

Nonetheless, recent technological advances, such as ribosome profiling (“Ribosome Profiling of Mouse Embryonic Stem Cells Reveals the Complexity and Dynamics of Mammalian Proteomes,” 2011), allowing for the global quantitative assessment of translation efficiency have unveiled a substantial lack of correlation between the transcriptome and protein levels (Y.-C. Wang et al., 2014). Notably during neurogenesis, the translational status of thousands of genes changes without matching variations in mRNA levels (Blair et al., 2017), most of them holding key functions in neural differentiation and function (Lins et al., 2022). These observations put forward that translational regulation has a major impact on the final output of gene expression. Moreover, the question how this control may be exerted on specific mRNA subsets has brought ribosomes as potential control elements into the limelight (Breznak et al., 2022).

In eukaryotes, a highly controlled and energy-consuming ribosome biosynthesis pathway ensures the correct assembly of this huge macromolecular complex made of RNA (rRNA) and proteins (RPs) (Baßler and Hurt, 2019; Khatter et al., 2015; Ramakrishnan, 2002). Beyond the core ribosome, a large number of associated factors have been identified in different organisms (Imami et al., 2018). Further complexity arises through post-translational modification of ribosomal proteins and the large number of different rRNA modifications (“Translational control through ribosome heterogeneity and functional specialization,” 2022). The two most abundant rRNA modifications are pseudouridines (ψ) and 2’-O-methylations (2’-O-me) (“Translational control through ribosome heterogeneity and functional specialization,” 2021). Both modifications are added to specific rRNA nucleotides by generic enzymes guided by small nucleolar RNAs (snoRNAs) *via* complementary base-pairing interactions. More specifically, ψ is installed by Dyskerin and guided by box H/ACA snoRNAs, and 2’-O-me executed by Fibrillarin in complex with box C/D snoRNAs (Steitz and Tycowski, 1995).

Despite the inherent complexity of the ribosome, investigation into the mechanisms by which translation is controlled has mainly focused on mRNA abundance, sequence, and secondary structure, as well as regulation by translation initiation and elongation factors (Genuth and Barna, 2018; Sternberg et al., 2009). However, over recent years, evidence has accumulated suggesting that ribosomes are not generic machines but come with a considerable amount of natural and pathologic variations. The sources for this diversity are manifold, and a direct consequence of the abovementioned complexity (“Translational control through ribosome heterogeneity and functional specialization,” 2022). As such, several studies have reported variation in the RP composition through the incorporation of RP paralogs or alterations in RP stoichiometry (Fusco et al., 2021; Gupta and Warner, 2014; Kondrashov et al., 2011; Shi et al., 2017; Slavov et al., 2015), and their post-translational modifications (Imami et al., 2018). Likewise, rRNA variation can stem from differential expression of variant rDNA alleles found in the multiple rDNA clusters present in mammalian cells (Fan et al., 2022; Parks et al., 2018), as well as from changes in the rRNA post-transcriptional modification profiles (Shi and Barna, 2015; Sloan et al., 2016; Xue and Barna, 2012). Furthermore, the observation in humans and mice that genetic defects of RPs, rRNA processing genes, or ribosome biogenesis factors may result in tissue-specific pathologies, termed ribosomopathies, indicates that the affected ribosomes play divergent roles in different cell types (Kampen et al., 2020; Khajuria et al., 2018). The establishment of ribosome heterogeneity has led to the hypothesis of functional ribosome specialization, where alternating core protein composition as well as protein or rRNA modifications could confer additional layers of regulation to the translation process by influencing translation speed and fidelity, or by promoting the translation of specific mRNA subsets (Norris et al., 2021; Shi and Barna, 2015; “Translational control through ribosome heterogeneity and functional specialization,” 2021).

Using RiboMeth-seq (RMS), a high-throughput sequencing-based method that allows for the simultaneous mapping and quantitative assessment of the 2’-O-me status of all rRNA residues (Birkedal et al., 2014; Krogh et al., 2017), we have previously observed that about a third of the 109 rRNA positions known to carry 2’-O-me in humans are fractionally methylated, i.e. that not all ribosomes of a given cell or tissue carry a modification at one of these positions (Jansson et al., 2021; Krogh et al., 2016). Moreover, these fractionally methylated sites in particular exhibit significant differences in their methylation degree between different cell types and conditions. These findings have been corroborated by studies demonstrating variation of the rRNA 2’-O-me-profile during normal development in zebrafish and mice (Hebras et al., 2020; Ramachandran et al., 2020), and in pathologies such as diffuse large B-cell lymphoma (Krogh et al., 2020) and breast cancer (Marcel et al., 2020). Together, the data suggest the existence of ribosome subtypes characterized by different 2’O-me modification patterns. Emerging experimental evidence support the notion of 2’-O-me sites facilitating ribosome specialization. For instance, we have recently shown that expression of Myc results in specific alterations of the ribosome 2’-O-me pattern in human cells, particularly at 18S:C174, which in turn impact translation of distinct mRNAs depending on their codon composition (Jansson et al., 2021).

Here, we aim to understand the importance of ribosomal 2’-O-me for cell fate establishment in early embryonic development and during neuronal specification. We show that the rRNA 2’-O-me profile undergoes significant and profound changes during mouse embryonic and postnatal brain development with dynamics at some positions being specific to certain brain regions. Tracing development back to germ layer specification, we demonstrate that the directed differentiation of human embryonic stem cells (hESCs) into the three embryonic germ layers trigger significant differentiation type-specific 2’-O-me dynamics. The importance of these dynamics is highlighted by our finding that the removal of a single, dynamic 2’-O-me modification push cell fate towards the neurectoderm. This is mediated through an altered translation of WNT signaling pathway members and differential association of the translational regulator Fragile X Mental Retardation Protein (FMRP) in the vicinity of the modulated 2’-O-me site.

Together, the data indicate that ribosomal RNA modification constitutes a previously unrecognized and essential regulatory mechanism in regulating mammalian gene expression and establishing cellular identity.

## Results

### Temporal and regional rRNA 2’-O-me dynamics during mouse brain development

Previously, we demonstrated the existence of 2’-O-me dynamics in cell culture models (Jansson et al., 2021). To investigate whether heterogeneity and dynamics of 2’-O-me exist *in vivo* during the transition from multipotent stem cells to differentiation, we focused on a developmental system with a tightly timed sequence of neurogenesis. We performed microdissection of mouse brain neocortex (CTX) during embryonic windows just prior to neurogenesis (E11), throughout neurogenesis (E12.5, E14, E15.5, and E17), and in the postnatal period after neurogenesis is complete (P0 and adult) (Fig. 1A). Subsequently, RMS quantification of the 109 known 2’O-me sites was performed on all samples in biological triplicates.

**Figure 1:**
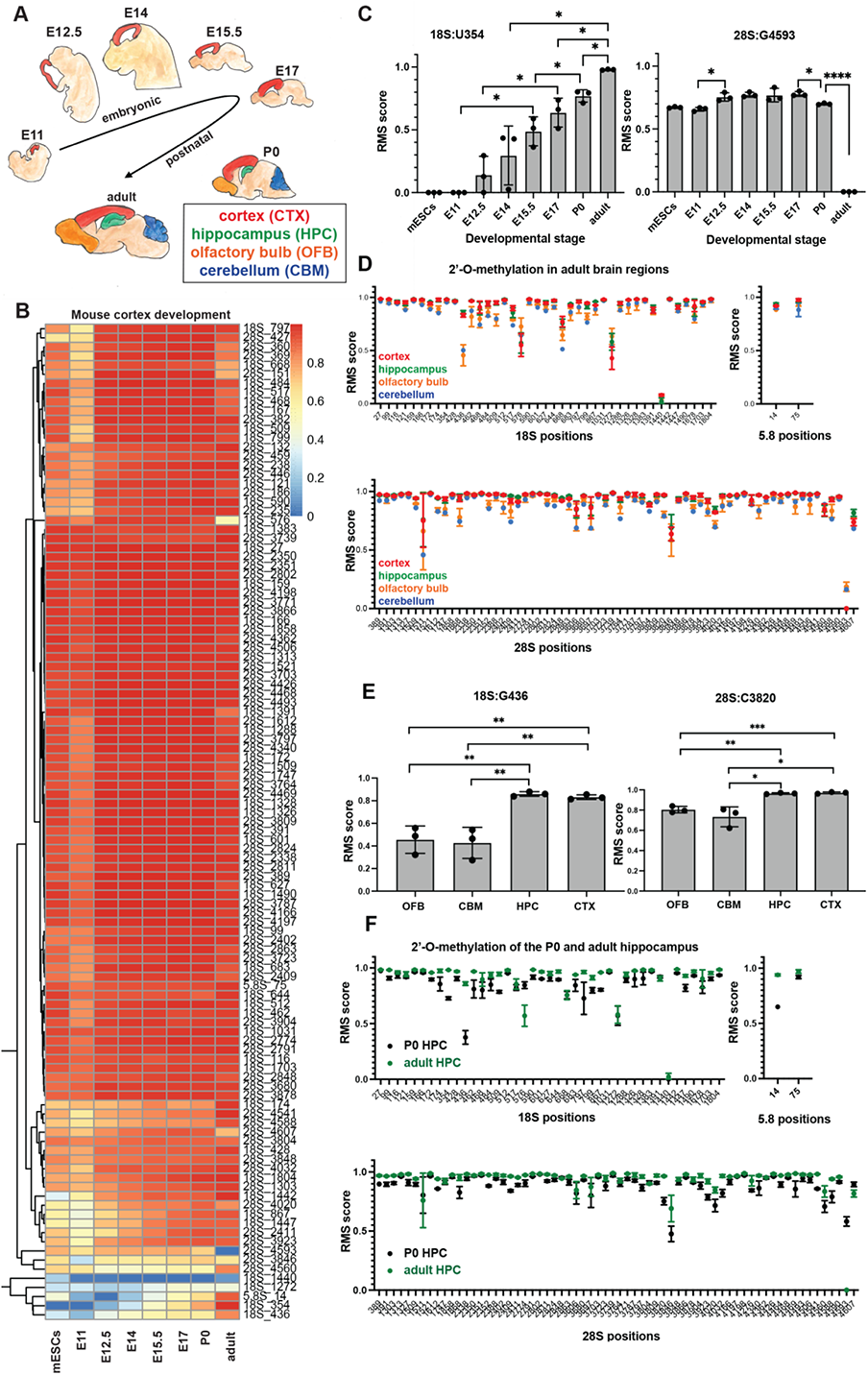
rRNA 2’-O-me dynamics in the developing mouse brain. A: Murine model system. Cortex (red) was obtained by dissection from 7 developmental stages ranging from E11 to adult. Hippocampus (green), olfactory bulb (yellow), and cerebellum (blue) were taken from neonates (P0) and adult. B: Heat-map depicting rRNA 2’-O-me levels in the developing mouse cortex and a mouse embryonic stem cell line measured by RiboMeth-seq. Columns represent developmental stages, rows all rRNA positions known to be potentially 2’-O-methylated from the 28S, 18S, and 5.8S rRNA. The color scale (from blue for low, to red for high) indicates the average RMS score from three biological replicates. C: RMS scores for two examples of rRNA positions displaying 2’-O-me dynamics during mouse cortex development, position 18S:U354 on the small subunit, and 28S:G4593 on the large subunit. Standard deviations refer to biological triplicates. ns: not significant. *: P≤0.05, **: P≤0.01, ***:P≤0.001, ****:P≤0.0001 (Welch’s unpaired t-test). D: Comparison of rRNA 2’O-me levels measured by RiboMeth-seq between four brain regions (cortex CTX, hippocampus HPC, olfactory bulb OFB, cerebellum CBM) from adult mice. Known methylated positions from the 28S, 18S, and 5.8S rRNA are depicted on separate graphs on the X-axis. The Y-axis corresponds to the average RMS score. Points represent mean RMS scores of *n* = 3 sequenced libraries from different animals. Error bars represent ± s.d. E: 2’-O-me levels at rRNA position 18S:G436 in different brain regions at the adult stage. Error bars represent ± s.d. of biological triplicates (points). ns: not significant. *: P≤0.05, **: P≤0.01, ***:P≤0.001, ****:P≤0.0001 (Welch’s unpaired t-test). F: As (D), for hippocampus of neonates (P0) (black) and adult mice (green).

We detect pronounced changes to rRNA 2’-O-me patterns over the course of cortex development (Fig. 1B). In accordance with previous observations (Jansson et al., 2021; Krogh et al., 2016), sub-stoichiometric methylation is detected at a subset of sites only, and significant changes in the degree of 2’-O-me, are seen at 43 sites (Supp. Table 1). Among the variable sites, most display an increase in 2’-O-me levels over the course of neocortex development. Some positions transit from undetectable to fully methylated (such as 18S:U354), while other sites display a late but substantial drop in methylation levels at the adult stage (for example 28S:G4593) (Fig. 1C). We observe hypo-methylation at embryonic stage (E11) when the neuroepithelium has yet to commit to a more restricted neural stem cell lineage at E12.5, giving rise to pyramidal neurons throughout subsequent embryonic stages. The 2’-O-me profile of mouse embryonic stem cells (mESC) cultured *in vitro* more closely resemble the multipotent E11 neuroepithelium (Fig. 1B). Interestingly, the majority of changes in the maturing neocortex are sequential and progressive over time, perhaps indicating a role for rRNA methylation dynamics in the stepwise acquisition of mature neuronal fate (Fig. 1B).

We next asked whether the neocortex rRNA methylation profile is aligned with other brain regions in the postnatal period. Towards that, we additionally micro-dissected hippocampus (HPC), cerebellum (CBM), and olfactory bulb (OFB) tissue from the same neonates and adult animals used for the cortex development analysis and performed RMS (Fig. 1D, Supp. Fig. 1A). We identified 9 positions with significant differences between at least two brain regions at the P0 stage (Supp. Table 2) and 8 positions in the adult (Supp. Table 3). Two examples in the adult, 18S:G436 and 28S:C3820, are shown in Fig. 1E. Both positions displayed higher methylation levels in the cortex and hippocampus compared to the cerebellum and the olfactory bulb. More generally, the 2’-O-me profiles of the cortex and hippocampus, and those of the cerebellum and olfactory bulb respectively form two separate clusters, consistent with their divergent neurodevelopmental origins (Supp. Fig. 1B). This indicates that different neuronal populations and/or distinct cellular compositions in different brain regions harbor differently methylated ribosome pools.

Subsequently, we extended our comparison to the 2’-O-me profiles of the same four brain regions between neonates (P0) and adult mice, revealing further marked differences (Fig. 1F, Supp. Fig. 1C, Supp. Table 4). Notably, the 2’-O-me level at 28S:G4593 drops substantially to almost zero in all brain regions only after birth (Supp. Table 4), demonstrating that dynamic changes in 2’-O-me continue to take place in the postnatal brain. Interestingly, 28S:G4593 is the only common postnatally dynamic position in all four regions, as both the number of significantly changing positions (7 in CTX, 3 in OFB, 6 in CBM, 9 in HPC), and their combination vary between the areas (Supp. Table 4). This reinforces the idea that the acquisition of regional identity is paralleled by the establishment of a specific combination of rRNA modifications and indicates a different composition of ribosome subtypes. Importantly, the 2’-O-me RMS values display a remarkable reproducibility between replicates, even though the samples come from independent animals. This suggests that the changes to rRNA 2’-O-me are tightly regulated during mouse brain development.

Together, these findings demonstrate that significant dynamics in rRNA 2’-O-me take place *in vivo* during mouse brain development, with modifications changing both across developmental time within the same brain region, and between distinct brain regions.

### Fate-specific 2’-O-me dynamics during human ESC differentiation

Our identification of dynamic rRNA 2’-O-me sites within the developmental transition from multipotency to differentiation in mouse neural tissue raised the question of whether 2’-O-me dynamics occur during earlier stages of stem cell committment. We therefore analyzed totipotent human ESC differentiation into the the three germ layers. For this purpose, we differentiated two human ESC lines (H9 and HUES4), as well as a human induced pluripotent stem cell (iPSC) line (KOLF2) into the three embryonic germ layers; endo-, meso-, and ectoderm (Fig. 2A). The ectoderm differentiation, in particular, consists of a stepwise restriction of pluripotency, first giving rise to early neural progenitors (eNPC), then to late neural progenitors (lNPC), and finally mature neurons (MN) (Fig. 2A). Appropriate generation of the desired cell types over the course of differentiation was confirmed by RT-qPCR analysis for a panel of pluripotency and germ layer markers (Supp. Fig. 2A, 2B) and immunohistochemistry (Supp. Fig. 2C). All three cell lines differentiated as expected, with the exception of ectoderm formation, where only the H9 cell line differentiated adequately and was thus used for further experiments (Supp. Fig. 2D). RMS was subsequently performed on all three cell lines in their pluripotent state and on their differentiated progeny (Fig. 2B, Supp. Fig. 3A). All three pluripotent cell lines showed very similar 2’-O-me profiles at both the pluripotent and differentiated stages, indicating that the observed dynamics were robust and reproducible (Supp. Fig. 3A).

**Figure 2:**
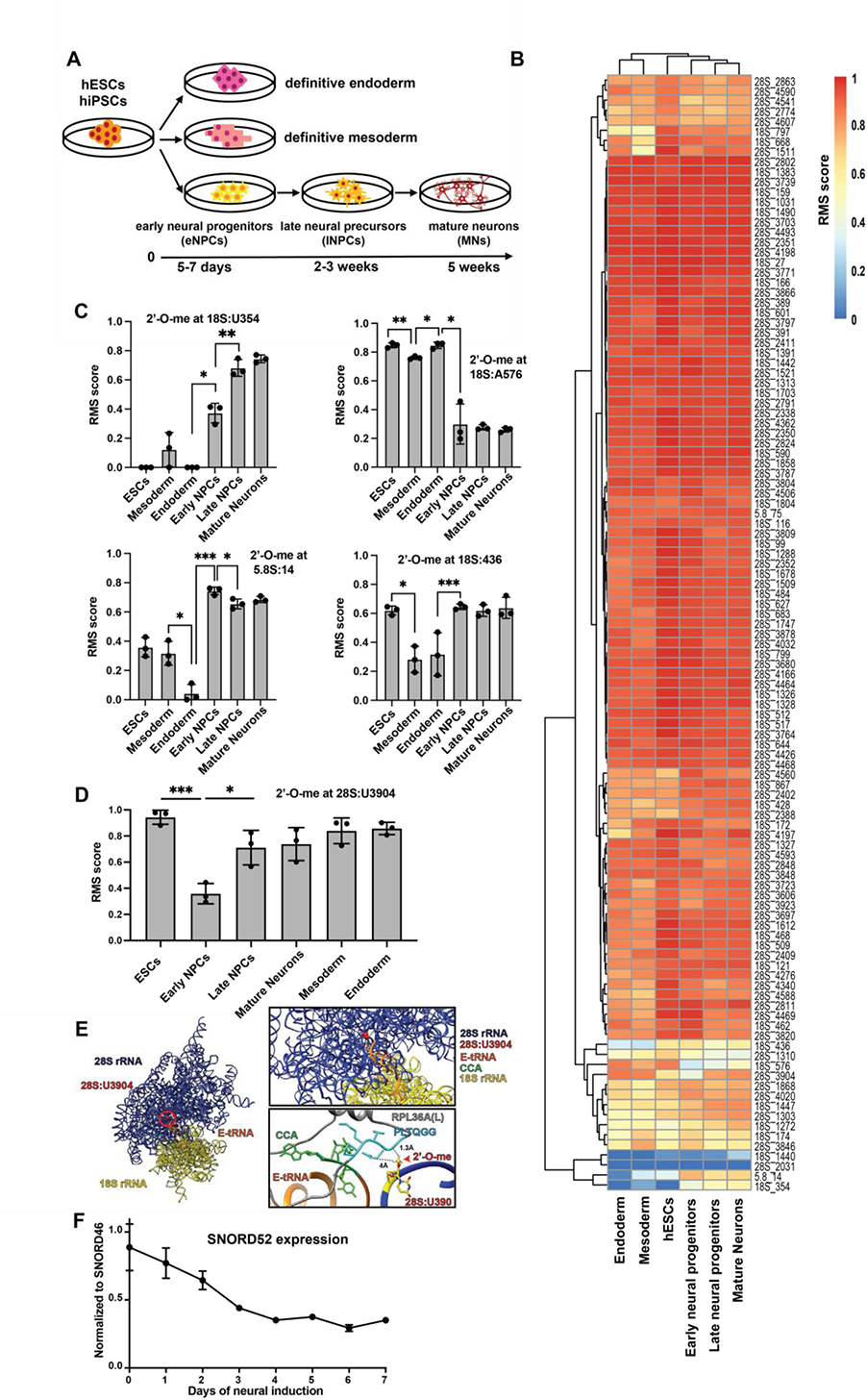
2’-O-me dynamics during directed differentiation of human ES cells. A: Human model system. Human embryonic stem cells (hESCs), H9 and HUES4, as well as human induced pluripotent stem cells (hiPSCs), KOLF2, were differentiated *in vitro* into the 3 embryonic germ layers endo-, meso-, and ectoderm. Neural differentiation represents ectoderm differentiation, subdivided into intermediate stages of cellular commitment, including early neural progenitor cells (eNPCs), late neural precursors (lNPCs), and mature neurons (MNs). The approximate time to reach the desired differentiation state is indicated. B: Heat-map showing RMS scores for H9 ^WT^ cells at the pluripotent stage (hESCs) and differentiated into endoderm, mesoderm, and ectoderm, respectively (columns). Ectoderm is further divided into early neural progenitor cells (eNPCs), late neural progenitor cells (lNPCs), and mature neurons (MNs). Rows correspond to all rRNA positions known to be potentially methylated from the 28S, 18S, and 5.8S rRNA. Methylation levels range from absent (RMS score = 0, blue) to full (RMS score = 1, red). C: Different types of rRNA 2’O-me dynamics during H9^WT^ differentiation into the 3 embryonic germ layers measured by RiboMeth-Seq. Cell stages are indicated on the X-axis, the Y-axis represents the fraction of rRNA molecules carrying a methylation at a certain position in these samples (RMS score). Standard deviations refer to biological triplicates. D: rRNA 2’-O-me dynamics at the large subunit position 28S:U3904 upon H9^WT^ differentiation into the three embryonic germ layers measured by Ribometh-Seq on biological triplicates. ns: not significant. *: P≤0.05, **: P≤0.01, ***:P≤0.001, ****:P≤0.0001 (Welch’s unpaired t-test). E: Left: localization of 28S:U3904 in the 3D structure of human rRNA. Blue: large subunit (28S). Yellow: small subunit (18S). Orange: tRNA. Red: 28S:U3904. Right, top: close-up of the human ribosome region harboring 28S:U3904, color code as (left). Right, bottom: 3D configuration of 2’-O-methylated 28S:U3904 (yellow), the CCA-portion of the E-tRNA (green/orange) and the conserved PLTQGG motif of ribosomal protein RPL36A(L) (cyan/grey). F: SNORD52 expression during the first week of neural induction of H9^WT^, assayed by RT-qPCR. Normalized to the housekeeping gene SNORD46. Error bars represent ± s.d.

Significant dynamics in 2’-O-me, at a subset of positions, were observed during the transition of stem cells into endo-, meso-, and ectoderm (Fig. 2B, Supp. Table 5). Strikingly, the combination of sites changing dynamically was differentiation type-specific, suggesting that different compositions of 2’-O-methylated ribosome subtypes are required for divergent differentiation processes (Fig. 2C, Supp. Table 5). For instance, position 18S:A576 was nearly fully methylated in pluripotency and maintained this level upon endo- and mesoderm differentiation, but methylation dropped significantly during ectoderm differentiation. In contrast, position 18S:U354 2’-O-me levels remained close to undetectable in all samples, except for ectoderm differentiation, where full methylation was gradually reached. As for position 18S:G436, the methylation level was stable during ectoderm differentiation as compared to pluripotent cells, but significantly decreased during endo- and mesoderm generation. Finally, position 5.8S:U14 displayed a different dynamics for each type of differentiation: the 2’-O-me level remains stable in mesoderm, decreases in endoderm, and increases in ectoderm differentiation (Fig. 2C). Moreover, the site-specific 2’-O-me dynamics observed *in vivo* during mouse brain development were largely recapitulated at the corresponding positions during human neurogenesis *in vitro*. As such, both models show a marked loss of methylation at position 18S:A576, and a substantial increase at position 18S:U354 over time (Fig. 1A-B and Fig. 2C).

Position 28S:U3904 sparked our interest, as it displayed intriguing 2’-O-me dynamics specifically upon neural differentiation: its methylation levels were high in hESCs and all three germ layers (RMS scores: 0.71-0.95), except for a transient drop to an RMS score of 0.36 at the early neural progenitor cell (eNPC) stage, the earliest cell fate commitment intermediate during ectoderm differentiation, often also referred to as “neurectoderm” (Fig. 2D). This observation prompted us to speculate that this transient decrease in 2’-O-me levels might be connected to, or even required for, epiblast-to-neurectoderm transition. 28S:U3904 is located in the immediate vicinity of the ribosome E site in the catalytic peptidyl transferase center and the ribosomal protein RPL36A(L) (Fig. 2E). 28S:U3904 2’-O-me is guided by SNORD52, and in line with the drop of methylation, the expression level of SNORD52 was also decreased upon neural induction, which recapitulates the epiblast-to-neurectoderm transition (in the present protocol, eNPCs emerge between days 7-10) (Fig. 2F).

Altogether, the data demonstrates significant germ layer-specific alterations to the rRNA 2’-O-me pattern and suggests that the ribosome population may be specifically altered to promote certain translational programs related to differentiation and cell fate establishment.

### Loss of 28S:U3904-me in hESCs shifts cell identity towards the neural fate

To investigate a potential causative link between specific 2’-O-me dynamics and cell fate in the transition from ESC to NPC in the ectoderm lineage, we manipulated the methylation levels of position 28S:U3904 by modulating the expression of the associated snoRNA guide. The genomic locus of SNORD52 is located in the third intron of a long noncoding RNA (lncRNA) of unknown function, *SNHG32* (Supp. Fig. 4A). This lncRNA hosts an additional snoRNA in its first exon, SNORD48, which guides the positioning of 2’-O-me at 28S:C1868 (Supp. Fig. 4A and 4B). With the striking exception of ectoderm differentiation, the 2’-O-me dynamics at these two positions were comparable (Supp. Fig. 4C). In addition, *SNHG32* was expressed at very low steady state levels in all cell types examined here, while SNORD48 levels are high, and SNORD52 displays moderate expression (Supp. Fig. 4D), thus arguing for a differential regulation of host genes and snoRNAs.

Using CRISPR-Cas9 editing (Supp. Fig. 4E), we excised SNORD52 from wild-type H9 hESCs (H9^WT^) and characterized two independent full knock-out clones (H9^52KO^) (Supp. Fig. 4F-G). Complete loss of methylation at 28S:U3904 was confirmed by RMS (Supp. Fig. 4H). Several CRISPR-negative clones (having undergone the exact same procedure as the H9^52KO^ clones but without a successful deletion) were analyzed in parallel and referred to as H9^CTRL^ (Supp. Fig. 4F).

Although cultured under the same stringent ESC conditions as the H9^WT^ cells and the H9^CTRL^ clones, the H9^52KO^ clones displayed marked morphological differences compared to the former. They exhibited features characteristic of eNPCs, such as small neurite outgrowths (Fig. 3A). Moreover, H9^52KO^ cells displayed lower proliferation rates compared to the wild-type (Supp. Fig. 5A). Strikingly, while both H9^52KO^ clones stained positive for the early neural transcription factor *PAX6*, expression of the pluripotency marker *OCT4* was markedly reduced in comparison to H9 wild type and H9^CTRL^ cells (Fig. 3B). Marker gene profiling by qRT-PCR of stemness and differentiation markers confirmed that the H9^52KO^ clones expressed decreased levels of several pluripotency marker genes, particularly *OCT4* and *NANOG*. In contrast, they displayed upregulation of a subset of ectoderm-specific markers, such as *NESTIN* and *SOX1* (Fig. 3C). These findings suggest that the loss of 28S:U3904 2’-O-me shifted the cellular identity of hESCs towards a neural fate. H9^52KO^ cells seem to adopt an identity of NPCs, where the levels of 28S:U3904 are naturally lowest (Fig. 2D). This would also explain the slight change in RMS score seen at positions 28S:A1310 and 28S:A3846 in the H9^52KO^ compared to H9^WT^ (Supp. Fig. 4H), given that these display a decrease and an increase in RMS score respectively during neurogenesis (Fig. 2B).

**Figure 3:**
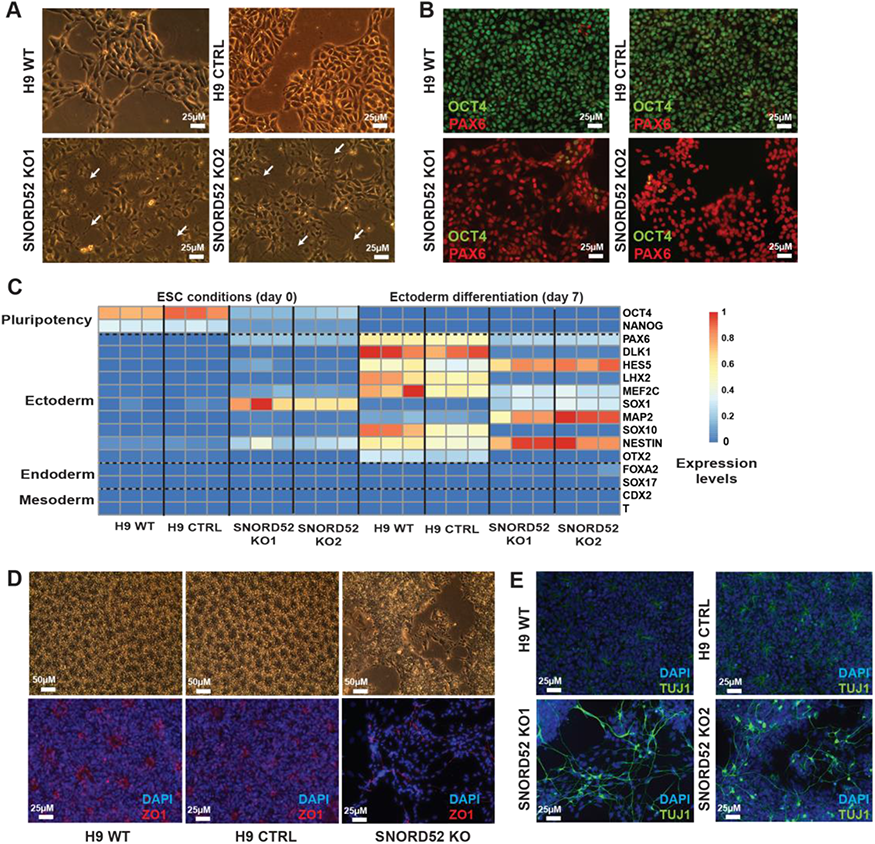
Loss of 2’-O-me at 28S:U3904 shifts cell identity from ESCs to NPCs under ES conditions and biases differentiation potential towards neurogenesis. A: Representative bright-field images of H9^WT^ ESCs (WT), CRISPR control H9^CTRL^ (CTRL), and two SNORD52 knock-out (H9^52KO^) clones (KO1, KO2) under hESC culture conditions. Magnification: 20x. White arrows indicate neuritic outgrowths. Representative images from n=3 experiments are shown. B: Immunofluorescence staining of H9^WT^, H9^CRTL^, and H9^52KO^ clones for the pluripotency transcription factor OCT4 (green) and the neurogenesis transcription factor PAX6 (red). The cells were grown under hESC culture conditions. Magnification 20x. Representative images from n=3 experiments are shown. C: Heat-map showing RT-qPCR assay of markers for pluripotency and differentiation into the 3 germ layers (rows) applied to H9^WT^, H9^CTRL^, and two H9^52KO^ clones as biological triplicates (columns) under hESC conditions (day 0), and at day 7 of ectoderm differentiation. Color scale indicates expression levels, from blue (low) to red (high). Values are normalized to GAPDH and to their respective expression ranges over the differentiation time-course. D: Bright-field and immunofluorescence images of H9^WT^, H9^CTRL^, and H9^52KO^ cells at day 7 of neural induction. Magnification 10x. Blue: DAPI. Red: ZO1, for visualization of neural rosette structures. Representative images from n=4 experiments are shown. E: Immunofluorescence staining at day 5 of neural differentiation of H9^WT^, H9^CTRL^, and two H9^52KO^ clones using nuclear staining (DAPI, blue) and an antibody against neuron-specific TUJ1 (β-tubulin III, green). Magnification: 20x. Representative images from n=3 experiments are shown.

To investigate the effect of manipulating 28S:U3904 2’O-me levels on differentiation and cell fate decision making, we subjected H9^WT^, H9^CTRL^, and H9^52KO^ cells to directed differentiation into the three embryonic germ layers. Upon ectoderm differentiation, H9^WT^ and H9^CTRL^ cells formed a dense monolayer patterned by neural rosettes, staining positive for ZO1 (Fig. 3D), typical for early forebrain progenitors, which is the expected default patterning in the absence of any added patterning factors (Pankratz et al., 2007). In contrast, H9^52KO^ cells grew less densely and did not form neural rosettes (Fig. 3D), but rapidly developed neurite-like extensions and networks, and strongly expressed the eNPC and lNPC marker NESTIN (Supp. Fig. 5B), as well as the later neural markers Tuj1 (β-tubulin III) (Fig. 3E) and MAP2 (Supp. Fig. 5B) well in advance compared to H9^WT^.

Marker gene expression profiling illustrated that on day 7 of ectoderm induction, H9^WT^, H9^CTRL^, and H9^52KO^ cell lines had effectively shut down pluripotency markers, and selectively up-regulated ectoderm-related genes (Fig. 3C, right). In contrast, H9^52KO^ cells displayed higher levels of *SOX1*, *NESTIN*, and *MAP2*, but lower levels of *SOX10*, *PAX6*, and *DLK1*. Interestingly, the H9^52KO^ cells displayed higher levels of *PAX6* at day 0, which remained relatively stable and seemed refractory to the transient upregulation of *PAX6* between days 3-5 as seen in H9^WT^ and H9^CTRL^ cells (Supp. Fig. 5C). Moreover, several genes failed entirely to be upregulated in the H9^52KO^ cells, including *OTX2*, *LHX2*, and *SOX10* (Supp. Fig. 5C). In addition, H9^52KO^ cells showed incomplete or failed endo- and mesoderm differentiation upon induction of these two germ layers and instead the cells formed atypical, tridimensional structures (Supp. Fig. 5D). Endoderm and mesoderm markers, like *CER1* or *T/BRACHYURY,* were either absent or displayed delayed upregulation (Supp. Fig. 5E and F). H9^52KO^-derived endoderm also displayed the aberrant expression of the ectoderm marker *SOX1* (Supp. Fig. 5E).

To assess if increased 2’-O-me levels at 28S:U3904 in hESCs would also impact cell identity, we overexpressed SNORD52 from an EGFP intron in H9^WT^ and characterized two H9^52OE^ clones. Stable expression of EGFP was verified at the ESC stage (Supp. Fig. 6A) and upon differentiation (Supp. Fig. 6B). As expected, we observed higher expression levels of SNORD52 at the ESC stage and throughout differentiation (Supp. Fig. 6C), and 2’-O-me levels at position 28S:U3904 were increased relative to the H9^WT^ under hESC conditions (Supp. Fig. 6D). Despite restoring 2’-O-me levels at 28S:U3904, re-introduction of SNORD52 into the H9^52KO^ cells did not revert them to the ESC stage, but rather induced growth arrest and terminal neural differentiation (Supp. Fig. 5G). Although the expression of germ layer-specific markers (Supp. Fig. 6E) suggest that H9^52OE^ cells are competent for differentiation into all three germ layers, the cells differed from H9^WT^ specifically for ectoderm differentiation. Notably, the cells did not form neural rosettes (Supp. Fig. 6F). Moreover, H9^52OE^ cells expressed some marker genes, such as *OTX2,* at higher levels than H9^WT^, which the H9^52KO^ cells failed to upregulate or expressed at low levels, (Supp. Fig. 6G).

These results indicate that abrogation of 2’-O-me at 28S:U3904 in hESCs suffices to drive the cells out of pluripotency and towards a neural cell fate, and potentially modulate the neurogenic potential.

### Loss of 2’-O-me at 28S:U3904 does not impact ribosome biogenesis

Installment of certain rRNA modifications, including 2’-O-me, are required for accurate ribosome biogenesis and assembly, generally through stabilizing local ribosome structure (Liang et al., 2007; Polikanov et al., 2015; Sloan et al., 2016). Northern blot analysis for the different rRNA intermediates revealed no imbalance indicative of ribosome biogenesis defect upon loss of 2’-O-me at 28S:U3904, although an overall reduction of processing intermediates was observed in both the H9^52KO^ cells and H9^WT^-derived eNPCs (H9^NPC^) compared to the H9^WT^ cells (Supp. Fig. 7A). This is consistent with several studies describing significant variations in both ribosome numbers per cell and overall translation during neural differentiation (Baser et al., 2019; Blair et al., 2017). Indeed, a peptide synthesis assessment *via* O-propargyl-puromycin (OPP) incorporation showed reduced global translation in H9^NPC^ and H9^52KO^ as compared to H9^WT^ cells, with the H9^52KO^ levels being similar to those of the H9^NPC^ (Supp. Fig. 7B).

Given that both ribosome and translation levels are comparable between H9^NPC^ and H9^52KO^ cells, we assume that the decrease in ribosome numbers that we observe in the H9^52KO^ cells is rather a consequence of the shift towards the neural fate than the result of a biogenesis defect, which fits previously reported observations (Chau et al., 2018).

### 28S:U3904 2’-O-me levels influence long-term neural cell identity

Following on from the clear bias towards neuroectoderm differentiation observed in H9^52KO^ cells, we proceeded to further characterize the neurogenic potential of H9^52KO^ cells by differentiating the cells into mature neurons, alongside H9^WT^ and H9^52OE^ -derived NPCs (H9^52OE-NPC^). Neural maturation was allowed for 50 days and was expected to produce primarily forebrain-type neurons and glia (Pankratz et al., 2007). In addition to morphology monitoring, the expression of a broad panel of brain region-specific markers was assessed by RT-qPCR at several time-points. Two weeks into the maturation phase, the H9^52KO^ cells consistently developed a distinct morphological phenotype as compared to the H9^WT^ and H9^52OE^ cells. While H9^WT^ and H9^52OE^ cells formed a homogeneous neural network, the H9^52KO^ cells grew first into a dense monolayer, then formed multiple circular cavities rimmed by thick borders with cilia-like cell protuberances (Fig. 4A). Gene expression analysis at the early neural induction phase (day 3 and 7) showed a marked suppression of forebrain-associated genes (*OTX2, OTX1, FEZF1, LHX2* and *SIX3*) and concomitant induction of hindbrain genes (*GBX2, HOXA1, HOXA2, EN1* and *OTP*), as well as induction of roof plate marker *PAX7* in the H9^52KO^ cells compared to the H9^WT^ cells (Fig. 4B).

**Figure 4:**
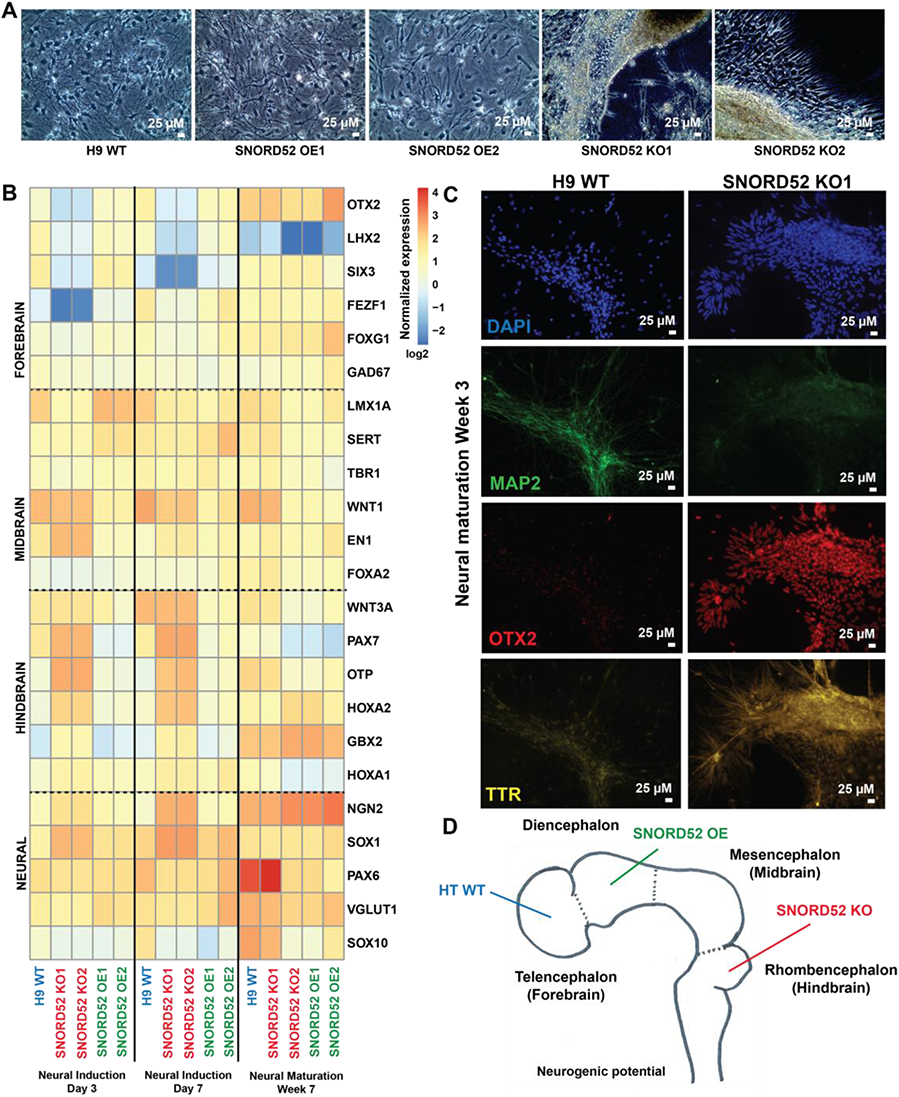
Manipulation of 28S:U3904 2’-O-me levels modifies the long-term neural differentiation potential of hESCs. A: Representative (of n=4) bright field images of H9^WT^, H9^52KO^, and H9^52OE^ cells at day 21 of neural maturation. Magnification: 20x. B: Heat-map based on RT-qPCR data for regional brain markers in H9^WT^, H9^52KO^, and H9^52OE^ cells at days 3 and 7 of early neural induction and after seven weeks of neural maturation. Genes are grouped by brain region. Values are normalized to GAPDH and reference values for a H9^WT^ cell line (also normalized to GAPDH), then transformed by log2. Color scale indicates expression levels, from blue (low) to red (high). C: Representative immunofluorescence images (of n=3) showing staining of H9^WT^ and H9^52KO^ cells after three weeks of neural maturation. Magnification: 20x. Blue: DAPI, green: MAP2, red: OTX2, yellow: TTR. D: Long-term differentiation potential of H9^WT^, H9^52KO^, and H9^52OE^ cells in terms of anterior-posterior brain regions.

Terminal maturation of the cultures further revealed the emergence of *TTR*+/*OTX2*+/*MAP2*-choroid plexus cells in the H9^52KO^ but not the H9^WT^ cultures, which is consistent with a H9^52KO^-induced shift in neural patterning from forebrain towards dorsal hindbrain fates (Rifes et al., 2020) (Fig. 4C).

H9^52OE^ cells in turn displayed an expression pattern similar to the one of H9^WT^ cells during early differentiation time points, but assumed a more posterior gene signature at the mature stage, indicating a diencephalon-like identity (Fig. 4B).

Altogether, the data indicate that the loss of 2’-O-me at 28S:3904 at the stem cell stage not only prompts the cells to shift to an NPC-like nature, but also programs them for terminal maturation into hindbrain neural cells, while the gain of 2’-O-me at 28S:3904 promotes differentiation into diencephalon cells (Fig. 4D).

### 2’-O-me at 28S:U3904 influences translation of mRNAs involved in WNT pathway

To investigate whether the methylation status of 28S:U3904 might influence the translation of specific mRNAs, and thus explain the early shift in cell identity in SNORD52 mutants, we performed ribosome profiling of H9^WT^ and H9^52KO^ cells under ESC culture conditions (Supp. Fig. 8A-D). Detected transcripts were categorized depending on their differential regulation in H9^52KO^ cells into those with significant differences in mRNA expression (transcription only); ribosome occupancy (translation only); both transcription and translation, opposite changes, or no change. The majority of changes (6615 transcripts) fell into the class of concordant changes in transcription and translation (Fig. 5A). Fitting previous observations, most transcriptionally upregulated genes in the H9^52KO^ cells were related to neural cell identity or function (Supp. Fig. 8E), thus confirming that the H9^52KO^ cells indeed shifted towards neuroectoderm. Interestingly, a subset of transcripts (1509) was significantly changed at the level of translation only (Fig. 5A). 708 transcripts displayed decreased translation (TL-DN) in the H9^52KO^ cells and gene ontology (GO) analysis revealed that these transcripts related primarily to translation and ribosome biogenesis, including ribosomal proteins (Supp. Fig. 8F). 801 transcripts were translationally upregulated (TL-UP) in the H9^52KO^ cells, and the most enriched, statistically significant GO category corresponded to genes related to the WNT signaling pathway (Fig. 5B). The WNT/β-catenin pathway plays a complex role in pluripotency and lineage commitment, sometimes taking on opposite functions depending on the spatiotemporal context (de Jaime-Soguero et al., 2018). On the one hand, WNT signaling is required for the induction and maintenance of stemness (de Jaime-Soguero et al., 2018), on the other hand, differentiation comes with the release of β-catenin from the cellular membrane (Sierra et al., 2018) and a surge of WNT transcriptional activity (Faunes et al., 2013; Sierra et al., 2018), which notably is required for neural induction (Mulligan and Cheyette, 2012; Otero et al., 2004) and rostro-caudal neural tube patterning (Fang et al., 2019; Rifes et al., 2020).

**Figure 5:**
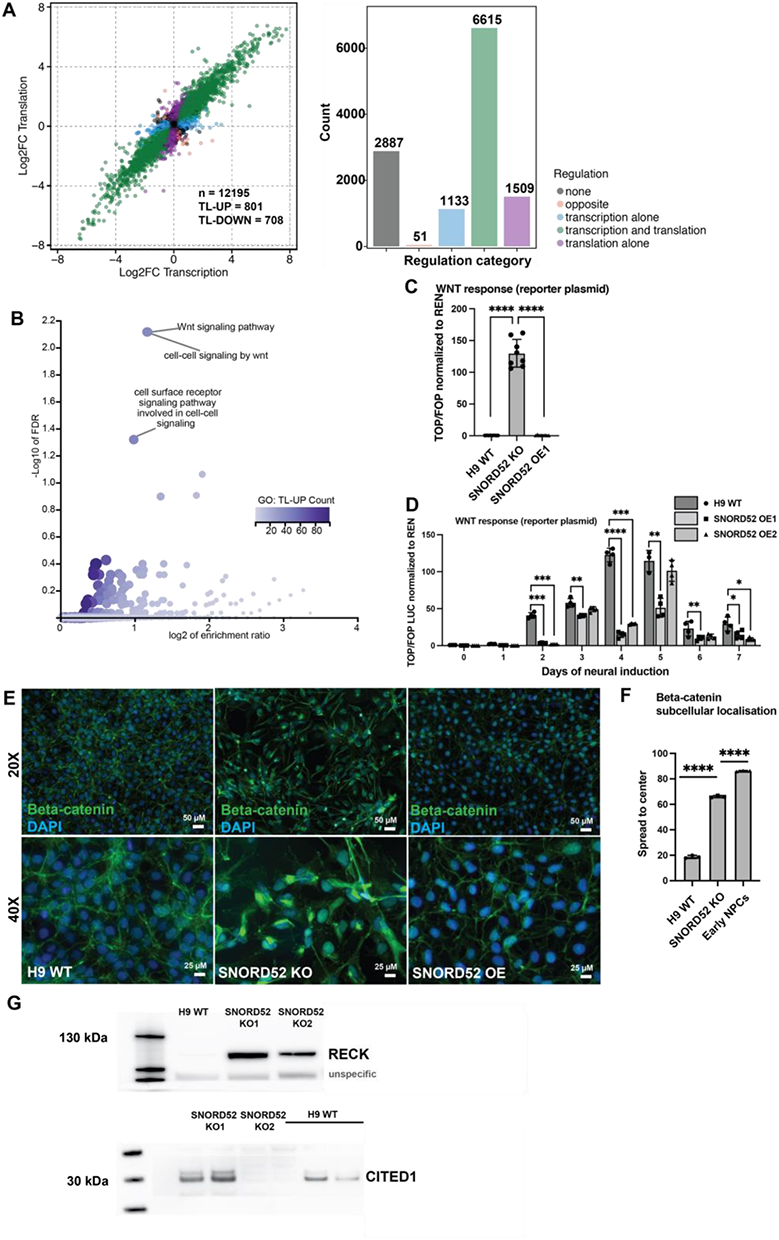
WNT pathway translation is repressed by 28S:U3904 methylation. A: Changes in mRNA expression and ribosome footprint levels from ribosome profiling data, comparing H9^WT^ and H9^52KO^ cells (n=3 libraries from individual cultures). Scatter plot (left) with mRNA transcripts colored to indicated regulation type. Number of sequenced transcripts analyzed given by *n*. Numbers of transcripts up- or down-regulated in H9^52KO^ relative to H9 WT cells in the translation alone set also shown (TL-UP, TL-DN, respectively). Histogram (right) giving the number of mRNA transcripts significantly regulated in each category (Benjamini–Hochberg *P*adj < 0.05). B: Gene ontology analysis of mRNA transcripts in TL-UP set. Biological process GO categories with FDR < 0.05 are labeled. Number of genes overlapping with each GO category indicated by the color-scale gradient. C: WNT activity measured by a TOP/FOP reporter assay in H9^WT^ and H9^52KO^ cells. Luciferase activity is used as readout for WNT activity and normalized to Renilla activity from a transfection control plasmid. The TOP plasmid contains WNT response elements, the FOP plasmid mutated versions of the latter. ns: not significant. *: P≤0.05, **: P≤0.01, ***:P≤0.001, ****:P≤0.0001 (Welch’s unpaired t-test). D: The same TOP/FOP reporter assay as in C with H9^WT^ and two H9^52OE^ clones over the course of the first seven days of neural induction. Average of three independent experiments. ns: not significant. *: P≤0.05, **: P≤0.01, ***:P≤0.001, ****:P≤0.0001 (Welch’s unpaired t-test). E: Immunofluorescence images of H9^WT^, H9^52KO^, and H9^52OE^ cells for beta-catenin (green) and nuclear staining (DAPI, blue). Top row magnification: 20x. Bottom row magnification: 40x. Representative images from n=3 biological replicates. F: Quantification of the subcellular localization of beta-catenin in H9^WT^, H9^52KO^, and early neural progenitors derived from H9^WT^ (H9^NPC^). ns: not significant. *: P≤0.05, **: P≤0.01, ***:P≤0.001, ****:P≤0.0001 (Welch’s unpaired t-test). G: Western Blot for RECK (left) and CITED1 (right) in H9^WT^ and H9^52KO^ cells under hESC conditions.

We assessed canonical WNT activity using the TOP/FOP luciferase reporter assay (Barolo, 2006). In line with the findings above, WNT activity was markedly higher in the H9^52KO^ cells compared to the H9^WT^ and ^H952OE^ under hESC conditions (Fig. 5C). Moreover, upon a neural induction time course, the induction of WNT activity was markedly lower in in the H9^52OE^ cells compared to the H9^WT^ (Fig. 5D). Release of β-catenin from the cell membrane, where it is found in a complex with the pluripotency factor OCT4, and its translocation to the cytoplasm and nucleus is an additional indicator of canonical WNT pathway activation (Faunes et al., 2013; Sierra et al., 2018). For this reason, we stained H9^WT^, H9^52KO^, and H9^52OE^ cells under ESC culture conditions for β-catenin. Levels were higher and β-catenin localization significantly more cytoplasmic and nuclear in the H9^52KO^ cells compared to the two other cell lines, where the staining was mainly detected at the cell membrane and was lowest in the H9^52OE^ cells, further supporting the finding that H9^52KO^ cells have activated canonical WNT signaling (Fig. 5E & F).

Western blotting furthermore confirmed the translational upregulation of WNT target genes in ^H952KO^ found by ribosome profiling, such as *RECK* and *CITED1 (Fig. 5G)*.

Hence, the data supports the notion that the methylation status of 28S:U3904 specifically impacts on the capacity of the ribosome population for translating specific mRNAs including those related to the WNT pathway.

### Ribosomes lacking 2’-O-me at 28S:U3904 display increased FMRP binding

To explore whether there are any differences in proteins associated with the ribosomes in H9^WT^, H9^52KO^, and H9^NPC^ cells, we purified ribosomes from 80S (monosome) and polysome fractions from each cell type and analyzed them by mass spectrometry (Supp. Table 7). In order to examine proteins most likely to be truly associated with the ribosome and remove the majority of contaminants present throughout the sucrose gradients, the detected proteins were filtered using a list of ribosome-interacting proteins derived from a previous study of two human cell lines (Imami et al., 2018) (Supp. Table 8, Supp. Table 9).

Overall, more proteins were found significantly associated with ribosomes from H9^WT^ compared to either H9^52KO^, or H9^NPC^ cells (Fig. 6A, Fig. 6B). These included many core ribosomal proteins (RPs) (Supp. Table 8), suggesting the samples from H9^WT^ cells contained more ribosomes in an equal amount of input material rather than indicating stoichiometric changes in so many individual RPs. This is consistent with downregulated ribosome biogenesis known to occur during differentiation (Breznak et al., 2022; R. Wang and Amoyel, 2022), in particular the strong translational repression of RPs, whose mRNAs are characterized by 5’ terminal oligopyrimidine (TOP) motifs (“Ribosome Profiling of Mouse Embryonic Stem Cells Reveals the Complexity and Dynamics of Mammalian Proteomes,” 2011), mediated by a decrease of mTOR activity (Blair et al., 2017; R. Wang and Amoyel, 2022).

**Figure 6:**
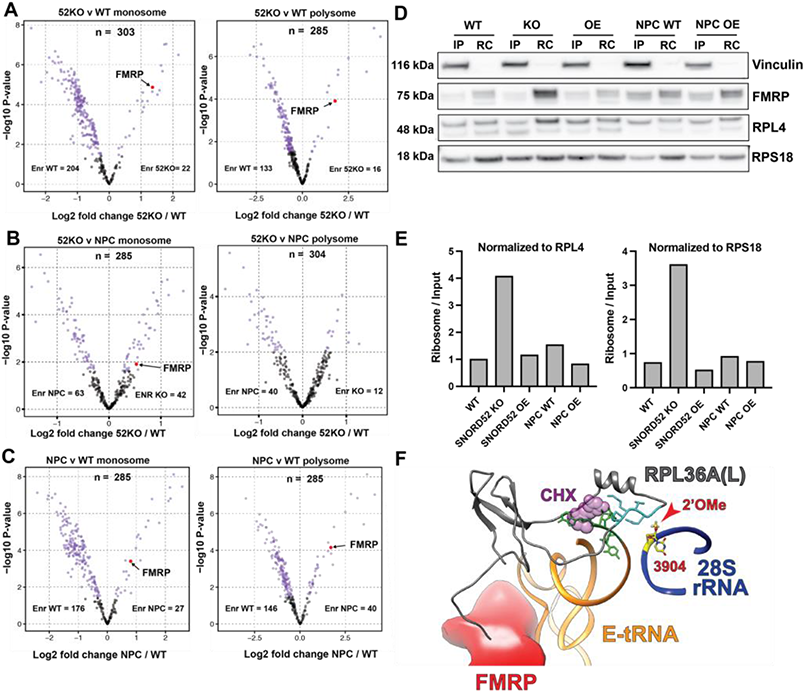
FMRP preferentially binds ribosomes lacking 28S:U3904 methylation. A: Protein abundance changes in H9^52KO^ (52KO) compared to H9^WT^ (WT) ribosome associated proteins in monosomes (left) and polysomes (right). Log 2 fold change in abundance (52KO/WT) and −log10 (*P* values) shown (*n* = 3 independent protein samples per condition). Those significantly changing (Benjamini–Hochberg *P*adj < 0.05) are colored (magenta). *n* denotes number of proteins analyzed. Numbers of proteins enriched in either sample is indicated (Enr WT, Enr 52KO). FMRP is highlighted in red. B: Protein abundance changes in H9^52KO^ (52KO) compared to H9^NPC^ (NPC) ribosome associated proteins in monosomes (left) and polysomes (right). log2 (abundance ratios) (52KO/WT) and −log10 (*P* values) shown (*n* = 3 independent protein samples per condition). Those significantly changing (Benjamini–Hochberg *P*adj < 0.05) are colored (magenta). *n* denotes number of proteins analyzed. Numbers of proteins enriched in either sample is indicated (Enr WT, Enr 52KO). FMRP is highlighted in red. C: Protein abundance changes in H9^NPC^ (NPC) compared to H9^WT^ (WT) ribosome associated proteins in monosomes (left) and polysomes (right). log2 (abundance ratios) (NPC/WT) and −log10 (*P* values) shown (*n* = 3 independent protein samples per condition). Those significantly changing (Benjamini–Hochberg *P*adj < 0.05) are colored (magenta). *n* denotes number of proteins analyzed. Numbers of proteins enriched in either sample is indicated (Enr NPC, Enr WT). FMRP is highlighted in red. D: Detection of total cellular and ribosome-bound FMRP protein by Western Blot in H9^WT^ (WT), H9^52KO^ (KO), H9^52OE^ (OE), H9^NPC^ (NPC WT), and H9^52OE^ -derived (NPC OE) cells. IP (input) = whole cell lysate, RC (ribosome cushion) = ribosomes purified by sucrose cushion. Vinculin serves as a control for appropriate separation of input and purified ribosomes, RPL4 and RPS18 for normalization to ribosome abundance. E: Quantification of the enrichment of FMRP in the ribosome-bound fraction based on (C). Bands were quantified, then subsequently the values for FMRP normalized to the corresponding ribosomal protein (RPL4 or RPS18), and finally the ratio of the ribosome-bound fraction over the input fraction calculated. See Supplementary Figure 9 for three more experimental repeats. F: 3D model of FMRP localization in the imminent vicinity of 28S:U3904, based on cryoEM data from drosophila. FMRP (red) is in direct contact with the E-site tRNA (orange) and the ribosomal protein RPL36A(L) (grey), which in turn lies within contact distance of 28S:U3904 (yellow, 28S rRNA in blue). The binding position of cycloheximide (CHX, violet) is additionally shown.

We focused on proteins enriched in samples from H9^52KO^ and H9^NPC^ cells compared to H9^WT^. Among these, we found FMRP (Fragile X Mental Retardation Protein) to be significantly more associated with ribosomes in H9^52KO^ (Fig. 6A), and H9^NPC^ (Fig. 6C), in both 80S and polysome fractions. Moreover, FMRP is also enriched in H9^52KO^ compared to H9^NPC^ in the 80S fraction (Fig. 6B). Despite having been shown to act at many levels of gene expression (Richter and X. Zhao, 2021), FMRP is best known for regulating translation of mRNAs involved in neurodevelopment (Li et al., 2020), leading to a crucial role in the regulation of the proliferation and cell fate of neural stem cells (Luo et al., 2010). FMRP can bind both to mRNA and to the ribosome, where it is assumed to bind within the intersubunit space and hamper the binding of tRNA and translation elongation factors (“Fragile X Mental Retardation Protein Regulates Translation by Binding Directly to the Ribosome,” 2014; Khandjian et al., 1996; Richter and X. Zhao, 2021).

Next, we sought to confirm the increased binding of FMRP to ribosomes devoid of 2’-O-me at 28S:U3904. We purified ribosomes from H9^WT^, H9^52KO^, H9^52OE^, and H9^NPC^ lines through sucrose cushions and probed for FMRP binding by western blot (Fig. 6D & Supp. Fig. 9). Two ribosomal proteins from different subunits (RPL4 and RPS18) were used for normalization to ribosome numbers. FMRP enrichment in the ribosome fraction was subsequently quantified as the ratio of ribosome-bound FMRP over total FMRP levels in the cell (input). We observe a strong enrichment of FMRP in the ribosome fraction of H9^52KO^ cells compared to all other cell lines (Fig. 6D and 6E, Supp. Fig. 9). Of note, WT^NPC^ cells display the second-highest levels of FMRP levels in the ribosome fraction, while in eNPCs derived from the H9^52OE^ cells (H9^52OE-NPC^) FMRP association was reduced, further consolidating the hypothesis that FMRP binding to the ribosome is favored by the absence of 28S:U3904 2’-O-me.

## Discussion

Correct development of complex mammalian organs, such as the brain, requires utmost spatiotemporal fine tuning of gene expression programs. Accordingly, numerous regulatory mechanisms have been revealed over the last decades. At the level of translation, focus has predominantly been on deciphering mechanisms governing translation initiation, elongation, and quality control. Yet, regulatory capacity through alterations of the rRNA modifications of the ribosome itself has remained understudied.

Over the last years, solid evidence has documented considerable ribosome heterogeneity in many organisms and recent studies have reported on ribosome specialization supporting translation of select mRNA populations, thus indicating a more profound role for the ribosome, or ribosome subtypes, in instigating specific translation programs (Genuth and Barna, 2018; Jansson et al., 2021; “Translational control through ribosome heterogeneity and functional specialization,” 2021). Although current technologies only allow for the analysis of ribosome composition or modification patterns in bulk across a tissue or cell culture, the evidence for ribosome heterogeneity strongly suggests the co-occurrence of multiple ribosome subtypes. Hence, dynamic changes in heterogeneity could be interpreted as an altered composition of functionally different ribosome subtypes acting in parallel within the same cell.

The current study significantly corroborates the contribution of rRNA 2’O-me variation to ribosome diversity and provides conceptually novel insight into the role of differential 2’-O-me in defining early stages of development and cell fate decision making during neurogenesis. We demonstrate profound rRNA 2’-O-me pattern differences between brain regions, which supports the hypothesis that ribosome diversification could contribute to the establishment of tissue identity. The observation that significant rRNA 2’-O-me dynamics take place *in vivo* during mouse brain development strongly implies biological importance. Most interestingly, 2’-O-me dynamics seem to persist into the postnatal period in the mouse brain. As the mammalian brain undergoes extensive restructuring of neural connections, also known as synaptic pruning, during the first weeks following birth, it is tempting to speculate that 2’-O-me and potentially other aspects contributing to ribosome heterogeneity are important for the fine-tuning of terminal cell identity in the neural network.

Tracking back to the early stages of development, we demonstrate that the directed differentiation of human ES cells into the three germ layers is paralleled by significant, robust, and germ layer-specific alterations to the 2’-O-me patterns of the ribosome population, suggesting a role for ribosome specialization in early development and cell fate decision making. The significance of these findings is further corroborated by the conservation of a number of dynamic 2’-O-me positions between the mouse *in vivo* and human *in vitro* neural differentiation models. This indicates an evolutionary conservation of 2’-O-me dynamics during neuronal development, at least among mammals.

Having established that ribosomal RNA 2’-O-me patterns consistently change during brain development and cell identity acquisition, we further demonstrated functionality by linking a single 2’-O-me position to a specific differentiation process and cell fate. Removal of the 2’-O-me at 28S:U3904 prompts hESCs to transition into neuroectoderm, despite being cultured under restrictive stemness conditions, and compromises their ability to differentiate into the two other germ layers. Furthermore, this influences the neurogenic potential of the cells by changing their regional identity upon long-term maturation towards a hindbrain nature. We did not observe changes in 2’-O-me levels at the murine 28S:U3904 locus in the brain development model. SNORD52 is duplicated in the mouse and the locus poorly conserved. Together, this could either signify an absence of dynamics in the mouse or indicate that they take place earlier or in a specific subset of neurons. The fact that the re-introduction of SNORD52 into the H9^52KO^ cells did not rescue the ESC phenotype is to be expected, given that the return to pluripotency, or “reprogramming”, is a complicated, low-efficiency process (Takahashi et al., 2007). In addition, as 28S:U3904 2’O-me increases again when NPCs differentiate into neural precursors and mature neurons, it is conceivable that expressing SNORD52, in an NPC-like context, rather promotes further progress down the neural cell fate than a return to the pluripotent state.

Using ribosome profiling, we identified a set of transcripts differing only at their level of translation following ablation of 2’-O-me at 28S:U3904. Most interestingly, gene ontology analysis strongly indicated a role for the canonical WNT pathway, and validation experiments confirmed active WNT signaling in the H9^52KO^ cells. The WNT pathway plays complex and multifaceted roles in cell identity, given that it operates in self-renewal and stemness as well as in lineage commitment (de Jaime-Soguero et al., 2018). De-repression or activation of WNT signaling either by inhibition of GSK3β (Shimojo et al., 2015) or activation of β-catenin signaling (Otero et al., 2004) facilitates the neural differentiation of hESCs. Furthermore, WNT participates in rostro-caudal organization of the neural tube (Rifes et al., 2020), axon guidance, as well as synapse development and activity (Mulligan and Cheyette, 2012). Regulation of WNT pathway members at the level of translation was functionally confirmed by the observation that H9^52KO^ cells display strongly upregulated WNT signaling, assume an NPC-like identity at the levels of morphology, gene expression, and differentiation potential, and finally are biased towards a hindbrain final cell fate, fitting the WNT gradient-governed pattern of the neural tube (Rifes et al., 2020).

Although we focused on the role of SNORD52 and 28S:U3904 2’O-me during neural differentiation, this does not exclude their implication in other developmental pathways. In a recent study, it was shown that inactivation of SNORD52 in hematopoietic stem cells increases subsequent erythroid differentiation (Nachmani et al., 2019). The hematopoietic lineages derive from the mesoderm, and indeed we observe a slight increase of the 2’-O-me levels at 28S:U3904 upon mesoderm differentiation, although we focused on early-stage mesoderm that has not yet undergone further specialization. Intriguingly, the WNT pathway also plays an important role in the specification of hematopoiesis (Sturgeon et al., 2014). It would be interesting to investigate the role(s) of the 28S:U3904 2’-O-me during the acquisition of non-neural cell identities.

An ongoing debate in the field relates to whether ribosome heterogeneity results in ribosome specialization (Ferretti and Karbstein, 2019; Mills and Green, 2017). An alternative to the ribosome specialization hypothesis proposes that varying the concentration of ribosomes may selectively impact different classes of mRNAs differently, without invoking specialized ribosome functions -the ribosome concentration hypothesis (Khajuria et al., 2018; Mills and Green, 2017). In contrast, other studies have demonstrated a specialized translation program resulting from, for instance, modulation of the ribosomal protein constituents (Shi et al., 2017) or rRNA modification patterns in the absence of changes to the ribosome concentration (Jansson et al., 2021; McMahon et al., 2019). The two views are not necessarily contradictory, and it is conceivable that some cases are regulated by a combination of specialized ribosomes and ribosome numbers, although disentangling the respective contribution of both mechanisms will be challenging. In the present study, loss of 2’-O-me at 28S:U3904 is accompanied by a drop in ribosome number in the H9^52KO^ cells. As such, we cannot fully rule out that part of the observed shift in cellular identity in our system is caused by a reduction in ribosome numbers during the transition from the ESC to the neuroectoderm state. Given that this reduction to similar levels is equally observed during the differentiation of WT ESCs to NPCs, it is likely that the phenomenon is rather another feature of the neural cell fate commitment than its triggering event.

To gain mechanistic insight into how 2’-O-methylation at 28S:U3904 can modulate translation, we purified ribosomes from H9^WT^, H9^52KO^, and H9^WT^ -derived neuroectoderm and analyzed their composition using mass spectrometry. This and subsequent validations revealed an increased binding of FMRP to ribosomes purified from H9^52KO^ and neuroectoderm derived from H9^WT^ cells. FMRP is a brain-enriched RNA-binding protein with a multitude of roles described related to translation, mRNA transport, splicing, and RNA stability (Hale et al., 2021; Richter and X. Zhao, 2021). Importantly, FMRP is also implicated in neurogenesis and neural cell fate (Li et al., 2020; Luo et al., 2010) and loss of FMRP leads to fragile X syndrome (Santoro et al., 2012). FMRP is generally considered a translational inhibitor and has been linked to the activation of WNT signaling through the translational repression of WNT inhibitors (Casingal et al., 2020; Luo et al., 2010). Such a scenario could link the enrichment of FMRP in H9^52KO^ cells with a subsequent activation of WNT signaling. FMRP has previously been found to interact with the 60S ribosomal subunit (Khandjian et al., 1996) and Cryo-EM structural analysis from Drosophila locates FMRP in the ribosomal intersubunit space in close vicinity of 28S:U3904 (“Fragile X Mental Retardation Protein Regulates Translation by Binding Directly to the Ribosome,” 2014) (Fig. 6E).

However, structural modeling predicts no direct interaction between FMRP and the 28S:U3904 rRNA residue, although this prediction is based on a low resolution Cryo-EM study in Drosophila. However, both FMRP and 28S:U3904 contact the E-site tRNA and the ribosomal protein RPL36A(L). RPL36A is eukaryote-specific (Kovacs et al., 2018), and one of the rare cases of a ribosomal protein with an active paralog in mammals, RPL36AL (Uechi et al., 2002). The paralogs differ only at the level of a single amino acid at position 38, a lysine in RPL36A and an arginine in RPL36AL. The ratio of ribosomes containing RPL36A over those bearing RPL36AL during development and normal homeostasis is unknown, but mutations in RPL36A(L) are linked with cycloheximide resistance (Klinge et al., 2011). In the ribosome, RPL36AL contacts both the CCA end of P-site bound tRNA and the translation termination factor eRF1 (Baouz et al., 2009; Hountondji et al., 2014). Most interestingly, RPL36A carries 7 and RPL36AL 6 monomethylated residues, among them Lys38, the one variable amino acid between the paralogs (Eustache et al., 2017), and the contact with the tRNA and eRF1 relies on the methylation of Lys35 (Eustache et al., 2017; Hountondji et al., 2012). In addition, there is evidence for fractional methylation of RPL36A(L) at positions Gln51 and Lys53, part of a highly conserved motif, which gave rise to speculations that such an extensive methylation pattern would likely be used for the regulation of translation or ribosome activity (Hountondji et al., 2012). Given that the 28S:U3904 is located less than 5Å proximity of the methylated GGQ motif of RPL36A(L), this could point towards an intricate interplay of both rRNA and RP heterogeneity through a methylation hotspot with direct effects on E-tRNA stability and FMRP binding.

Altogether, our findings reinforce the idea that the ribosome itself is a direct regulator of translation and demonstrates that modulation of the ribosome through alterations in the rRNA 2’O-me modification pattern contributes to directed differentiation and cell fate decision making during early development.

## Acknowledgements

The Lund lab is supported by grants from the Danish Council for Independent Research (Sapere Aude program 418 3-00179B); the Novo Nordisk Foundation (NNF18OC0030656, NNF 0071919); the Lundbeck Foundation (R198-2015-174), and the Danish Cancer Society (R204-A12532). Furthermore, this project has received funding from the European Union’s Horizon 2020 research and innovation program under the Marie Skłodowska-Curie grant agreement n° 801481 and The VILLUM Experiment Programme under the grant n° 17544. The authors would like to thank Bettina Mentz, Patrice Ménard, Anna Fossum, and Elin Josefina Pietras for their technical help.

## Author contributions

S.J.H. designed most experiments, collected data and drafted the article. M.D.J. carried out ribosome profiling and data interpretation, and mass spectrometry data analysis. K.A. was in charge of bioinformatic data analysis. M.L.K. provided mouse brain samples and insight into brain development and ribosome structure. K.L.A. carried out mass spectrometry data collection. Z.C. and A.N. assisted with qPCR data collection and data interpretation. M.F and D.M.S. were involved in establishing cell lines. D.M.G., F.S., and D.T provided critical input on the manuscript. N.K. contributed ribosome biogenesis-related experiments. H.N and A.K. provided advice on data interpretation. A.H.L. and M.D.J. assisted with work conception, data interpretation and drafting of the manuscript. All authors commented on the manuscript.

## Competing interests

The authors declare no competing interests.

## Supplementary Figures

**Supplementary Figure 1:**
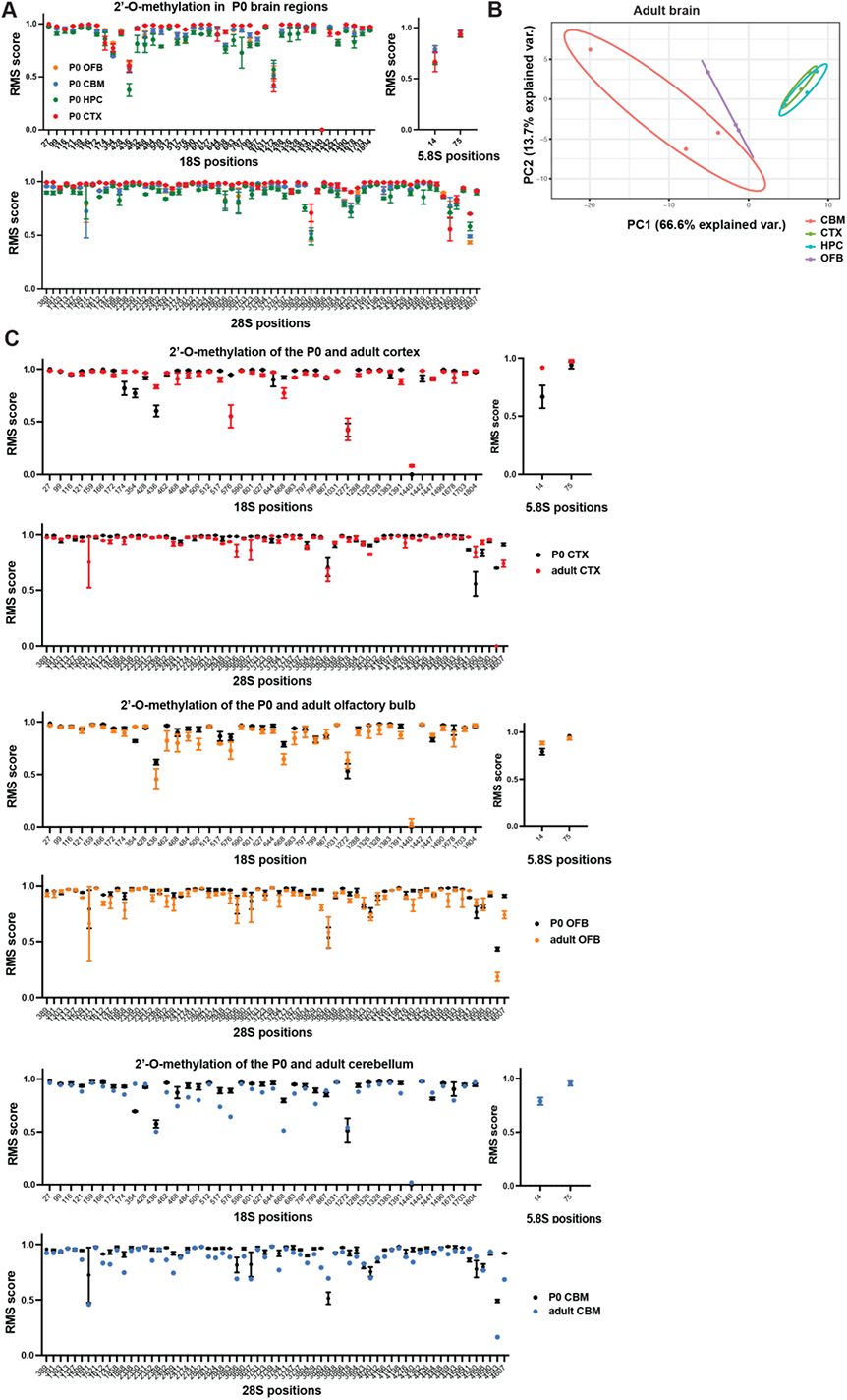
Additional RMS analyses of adult and neonate mouse brain regions. A: Comparison of rRNA 2’O-me levels measured by RiboMeth-seq between four brain regions (cortex CTX, hippocampus HPC, olfactory bulb OFB, cerebellum CBM) from neonate mice (P0). Known methylated positions from the 28S, 18S, and 5.8S rRNA are depicted on separate graphs on the X-axis. The Y-axis corresponds to the average RMS score. Points represent mean RMS scores of *n* = 3 sequenced libraries from different animals. Error bars represent ± s.d. B: PCA analysis of RMS data from four adult mouse brain regions: cerebellum (CBM), cortex (CTX), hippocampus (HPC), and olfactory bulb (OFB). 3 libraries (biological replicates) per brain region. C: Pairwise comparison of rRNA 2’-O-me levels between adult and neonates (P0) for cortex (CTX), olfactory bulb (OFB), and cerebellum (CBM). RiboMeth-seq (RMS) scores representing fraction of 2’-O-methylation at each potentially methylated site in 18S, 28S, and 5.8S rRNAs present in total RNA purified from the indicated brain regions. Nucleotide positions in respective rRNAs are indicated. Points represent mean RMS scores of *n* = 3 sequenced libraries from different animals. Error bars represent ± s.d

**Supplementary Figure 2:**
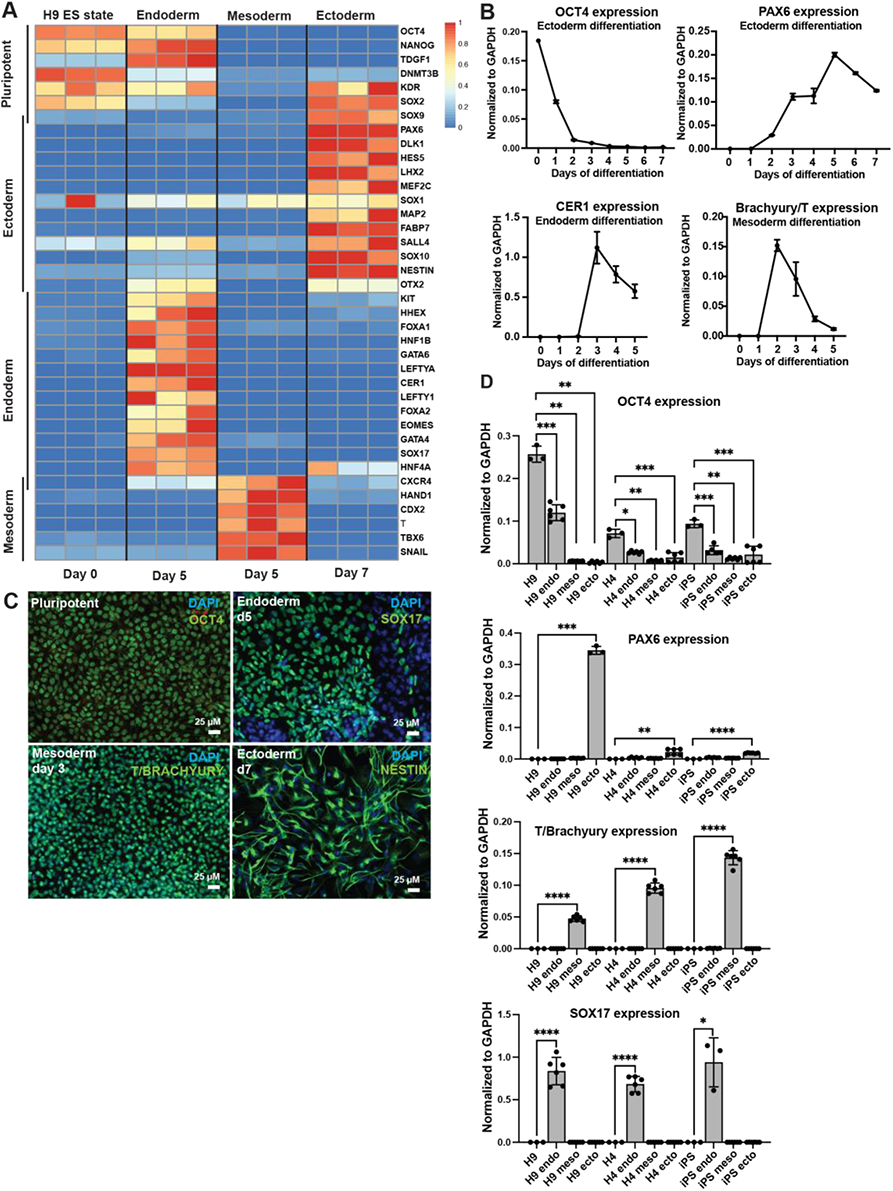
Confirmation of efficient directed *in vitro* differentiation of hESCs into the three embryonic germ layers. A: Heat-map based on RT-qPCR data confirming expression of known marker genes (rows) for pluripotent H9^WT^ hESCs and their differentiated progeny (columns) on the final day of differentiation (day 7 for ectoderm, day 5 for endoderm and mesoderm). Three independent experiments are plotted as separate columns per differentiation type. Blue corresponds to low expression levels, red to high levels. The genes are grouped by cell type (stem cells or germ layer). Values are normalized to GAPDH and to their respective expression ranges over the differentiation time-course. B: Time-course of mRNA expression normalized to GAPDH for representative marker genes during differentiation of H9^WT^ hESCs into the three germ layers measured by RT-qPCR. OCT4: pluripotency marker, PAX6: ectoderm marker, CER1: endoderm marker, T/Brachyury: mesoderm marker. Error bars indicate ±SD of n=3 biological replicates. C: Immunofluorescence staining for pluripotency and appropriate differentiation into the three germ layers in H9^WT^. Blue: DAPI. Green: germ line-specific differentiation markers. OCT 4 (pluripotency) under hESCs culture conditions, SOX17 at day 6 of endoderm differentiation, T/BRACHYURY at day 3 of mesoderm differentiation, and NESTIN at day 7 of ectoderm differentiation. Magnification: 20x. Representative images from n=3 experiments are shown. D: Comparison of the differentiation potential into the three embryonic germ layers of different pluripotent cell lines (H9, HUES4, and Kolf2 iPS cells) by RT-qPCR. Values are normalized to GAPDH. Error bars indicate ±SD from n=3 independent experiments.

**Supplementary Figure 3:**
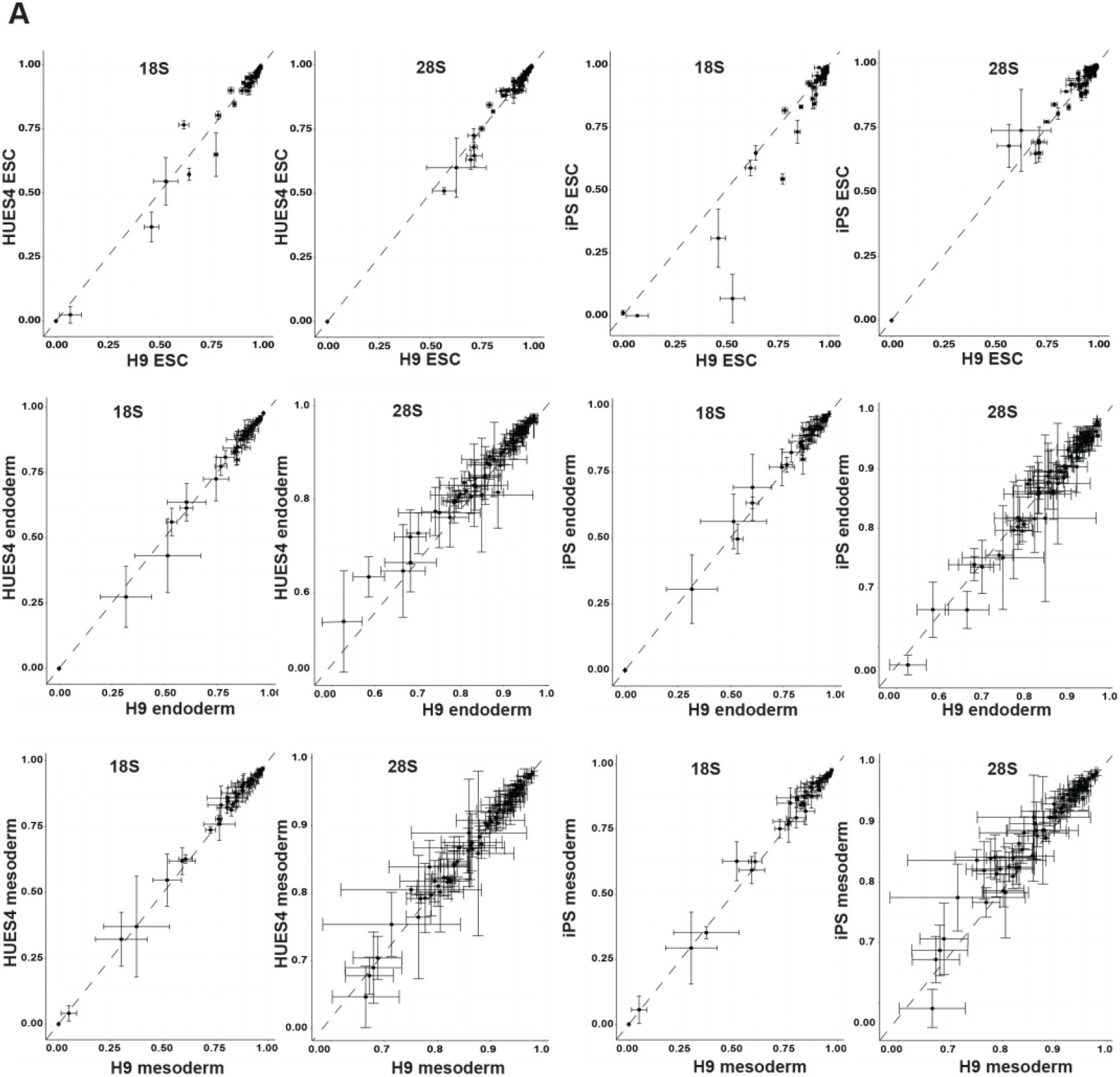
Differentiation-related rRNA 2’-O-me dynamics are reproducible between different pluripotent cell lines. A: Two by two correlation between cell lines (H9, HUES4, Kolf2 iPS) for RMS values at the pluripotent stage or after differentiation into endoderm and mesoderm for all known rRNA positions to be potentially 2’-O-methylated. Small subunit (18S) and large subunit (28S) positions are plotted in separate graphs. Error bars indicate ±SD from n=3 independent experiments.

**Supplementary Figure 4:**
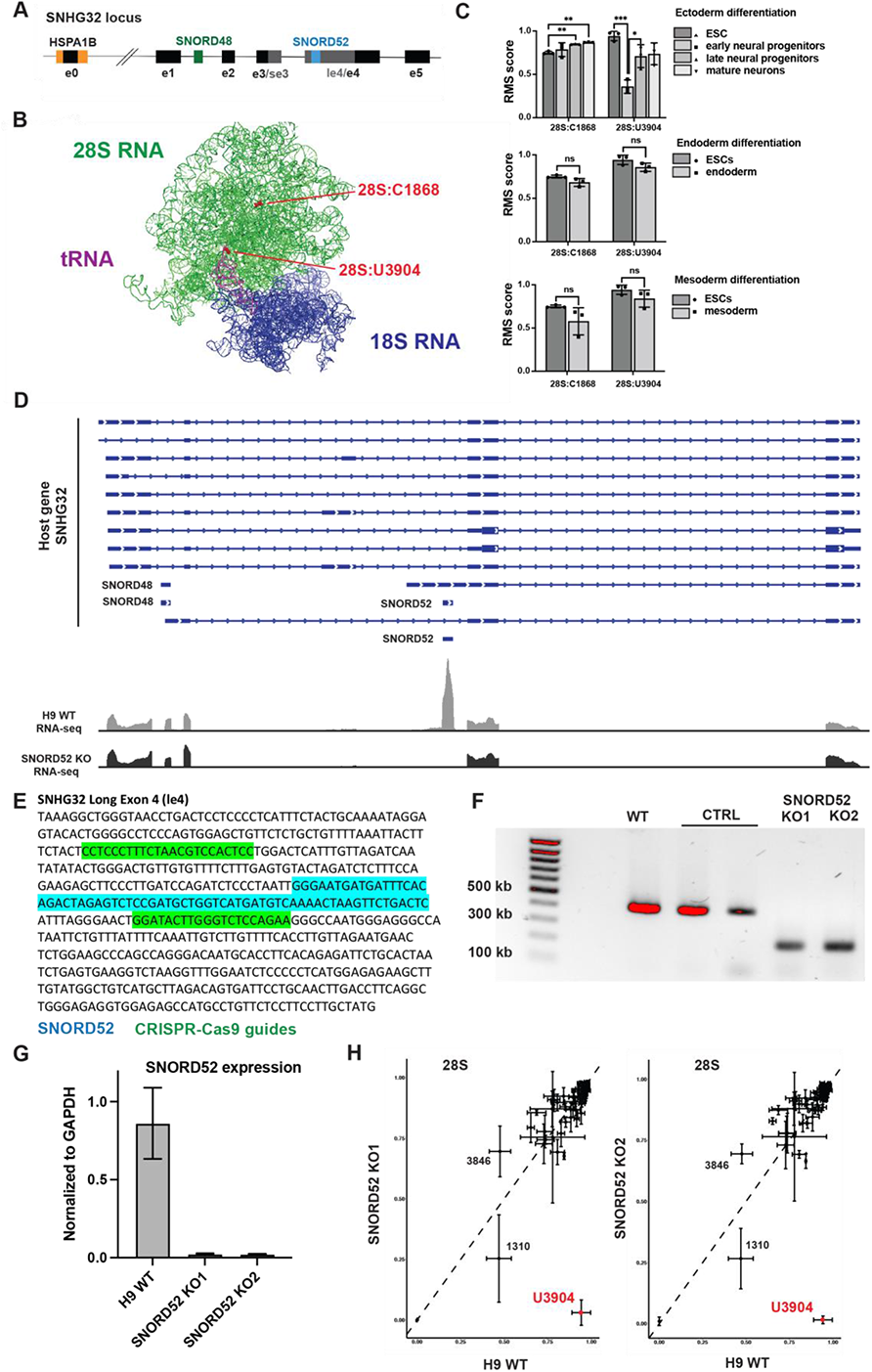
Characteristics of the SNORD52 locus and generation of SNORD52 knock-out clones by CRISPR/Cas9. A: Genomic locus of SNORD52, SNORD48, and their host gene SNHG32. e0-e5: SNHF32 exons. se3: short exon 3. le4: long exon 4. HSPA1B: Heat Shock Protein Family A Member 1B gene. B: Localization of 28S:U3904 (guided by SNORD52) and 28S:C1868 (guided by SNORD48) in the human rRNA 3D structure (PyMol graphic). Large subunit (28S) in green, small subunit (18S) in blue, 5.8S in turquoise, tRNA in magenta and methylation sites in red. C: Respective rRNA 2’O-me dynamics of 28S:C1868 and 28S:U3904 during the differentiation into the three embryonic germ layers. Error bars indicate ±SD from n=3 independent experiments. D: RNAseq tracks for the SNGH32 gene in H9^WT^ (H9 WT) and H9^52KO^ (SNORD52 KO) cells under hESC culture conditions. E: Sequence of SNORD52 preceding exon 4 of SNHG32, as well as the location and sequence of the CRISPR/Cas9 guides used to excise SNORD52 to generate the H9^52KO^ clones. F: PCR products for the region spanning the SNORD52 genomic region run on agarose gel to confirm the deletion of SNORD52 in the two KO clones. Control (H9^CTRL^) clones have undergone transfection with the CRISPR/Cas9 guides, but no deletion has taken place and the SNORD52 locus is intact. G: Confirmation by RT-qPCR of the absence of SNORD52 expression in the two H9^52KO^ clones compared to H9^WT^ cells. Values were normalized to GAPDH. Error bars indicate ±SD of n=3 technical triplicates. H: Deletion of SNORD52 fully removes 2’-O-methylation at 28S:U3904. RMS values for all known 2’O-me positions at the large subunit (28S) compared between H9^WT^ and the two H9^52KO^ clones. 28S:U3904 is highlighted in red. Error bars indicate ±SD of n=3 sequenced libraries from individual cell cultures.

**Supplementary Figure 5:**
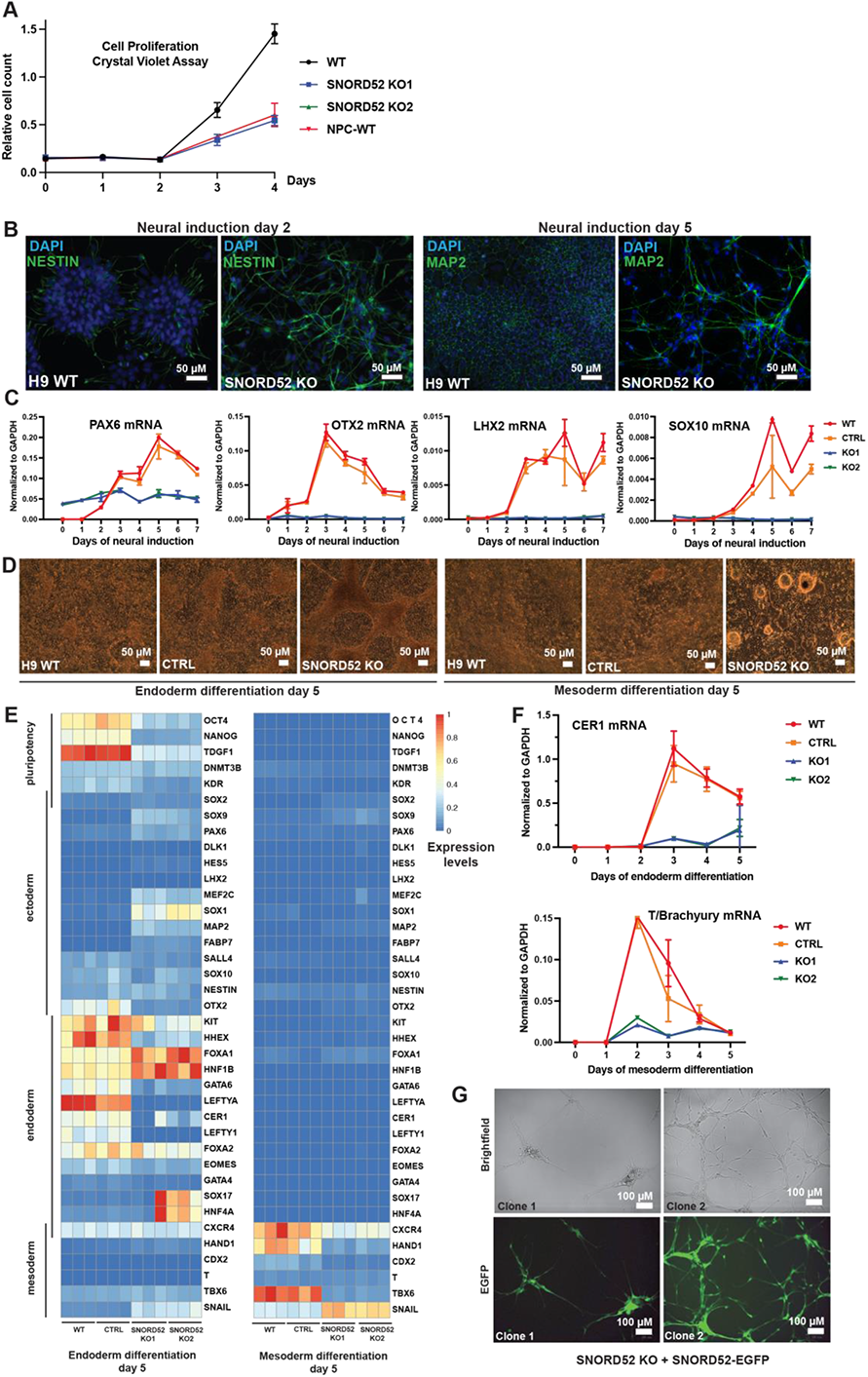
SNORD52 deletion correlates with failure to up-regulate certain neural markers and impedes differentiation into endoderm and mesoderm. A: Proliferation assay by crystal violet for H9^WT^, H9^52KO^ clones, and H9^WT^-derived neural progenitor cells (H9^NPC^). Error bars indicate ±SD of n=3 replicates. B: Immunofluorescence staining for ectoderm markers NESTIN and MAP2 in H9^WT^ and H9^52KO^ cells at day 2 (left) and 5 (right) of neural induction. Blue: DAPI. Green: NESTIN or MAP2. Magnification: 20x. Representative images from n=3 experiments are shown. C: Time-course of ectoderm marker gene (*PAX6, OTX2, LHX2, SOX10*) expression normalized to GAPDH during neural induction on H9^WT^ (WT), H9^CTRL^ (CTRL), and H9^52KO^ clones (KO1 and KO2) cells by RT-qCPR. Error bars indicate ±SD of n=3 biological replicates. D: Bright-field microscopy images of H9^WT^, H9^CTRL^, and H9^52KO^ cultures at day 5 of endoderm (left) and mesoderm (right) differentiation. Magnification: 10x. Representative images from n=3 experiments are shown. E: Heat-map based on RT-qPCR data for pluripotency and differentiation markers (columns) for H9^WT^, H9^CTRL^, and H9^52KO^ cells at day 5 of endoderm (left) or mesoderm (right) differentiation. Three independent experiments are plotted as separate columns per differentiation type. Values are normalized to GAPDH and subsequently to the range of all values per primer pair throughout the entire differentiation time-course (day 0 – day 5). Blue corresponds to low expression levels, red to high levels. The genes are grouped by cell type (stem cells or germ layer). F: Time-course of endoderm marker *CER1* and mesoderm marker *T/Brachyury* mRNA expression normalized to GAPDH during the differentiation of H9^WT^, H9^CTRL^, and H9^52KO^ cells into endoderm and mesoderm respectively assayed by RT-qPCR. Error bars indicate ±SD of n=3 biological replicates. G: SNORD52 was re-inserted into H9^52KO^ cells by targeting the AAVS1 locus with a 2-exon EGFP construct harboring SNORD52 in its intron. Cells did not revert to the ESC stage but differentiated terminally into neural-like cells while expressing high levels of the EGFP-SNORD52 construct. Two representative clones are shown. Top: bright-field. Bottom: EGFP. Magnification: 40x.

**Supplementary Figure 6:**
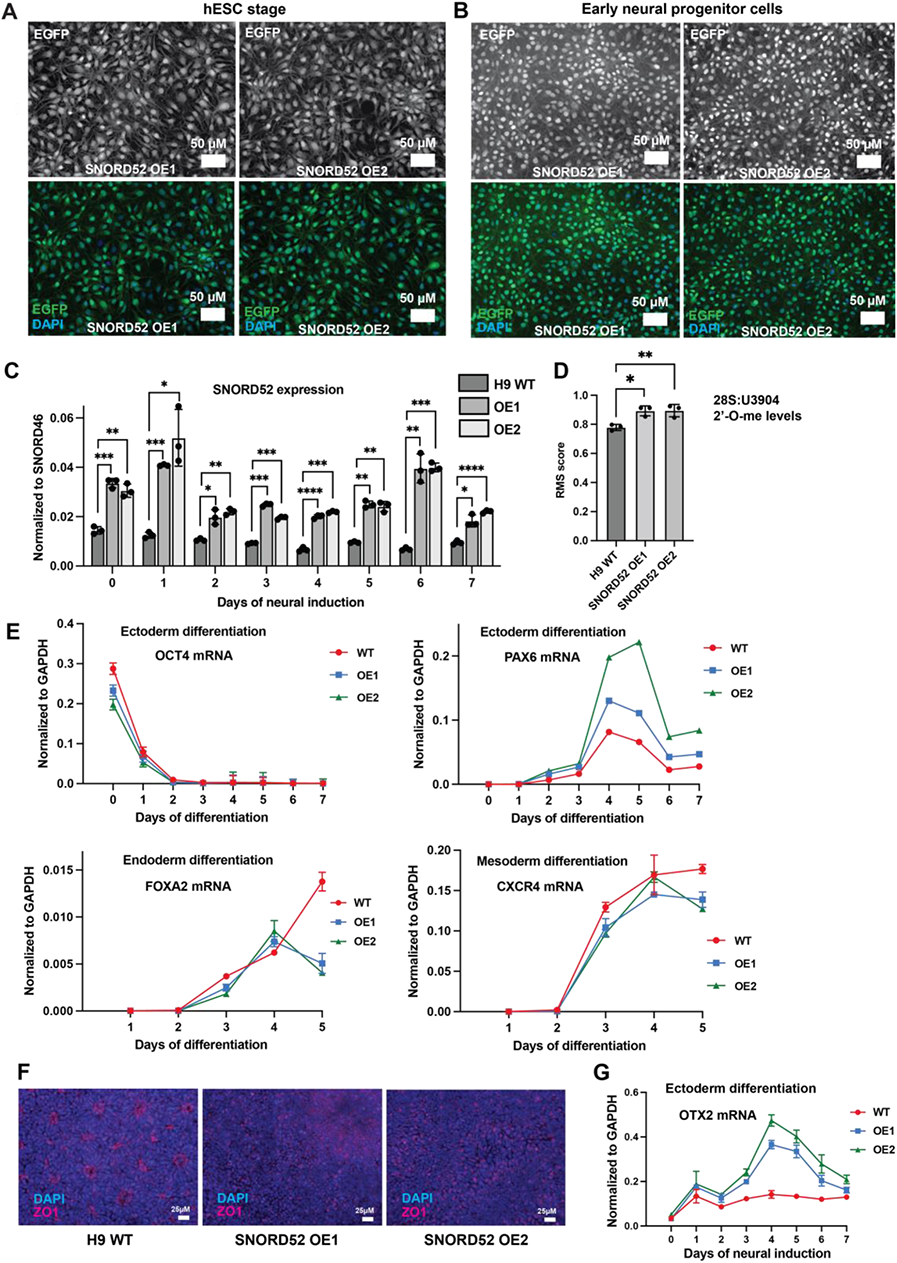
Generation and characterization of SNORD52 constitutive overexpression cell lines. A: Fluorescence images of two H9-derived cell lines with constitutive SNORD52 overexpression (H9^52OE^, OE1 and OE2) by targeting the AAVS1 locus with a 2-exon EGFP construct harboring SNORD52 in the intron. Top panels show EGFP, bottom panels EGFP and DAPI. Blue: DAPI. Green: EGFP. Magnification: 20x. B: Fluorescence images of H9^52OE^ cells following differentiation. H9^52OE^-derived early neural progenitor cells (H9^52OE-eNPC^) are shown. Top panels show EGFP, bottom panels EGFP and DAPI. Blue: DAPI. Green: EGFP. Magnification: 20x. C: Constitutive expression of the EGFP-SNORD52 construct achieves overexpression of SNORD52 throughout ectoderm differentiation. RT-qPCR data from neural induction in H9^WT^ (WT) and H9^52OE^ cell lines (OE1 and OE2). Normalization to SNORD46. Error bars indicate ±SD of n=3 technical triplicates. D: 2’-O-me at LSU 3904 in H9^52OE^ cells (OE1 and OE2) under hESC culture conditions. RMS scores shown. Error bars indicate ±SD of n=3 sequenced libraries from individual cell cultures. E: Pluripotency (*OCT4*) and germ-layer specific marker gene expression (*PAX6, FOXA2, CXCR4*) during differentiation in H9^WT^ (WT) and H9^52OE^ cells (OE1 and OE2), measured by RT-qPCR normalized to GAPDH. Error bars indicate ±SD of n=3 technical replicates. F: Representative fluorescence images of staining for ZO1 (red) and DAPI (blue) in order to highlight neural rosette structures at day 7 of ectoderm differentiation in H9^WT^ and H9^52OE^ cells. G: Expression of *OTX2* over neural induction in H9^WT^ (WT) and H9^52OE^ (OE1 and OE2) cells measured by RT-qPCR. Values normalized to GAPDH. Error bars indicate ±SD of n=3 independent experiments.

**Supplementary Figure 7:**
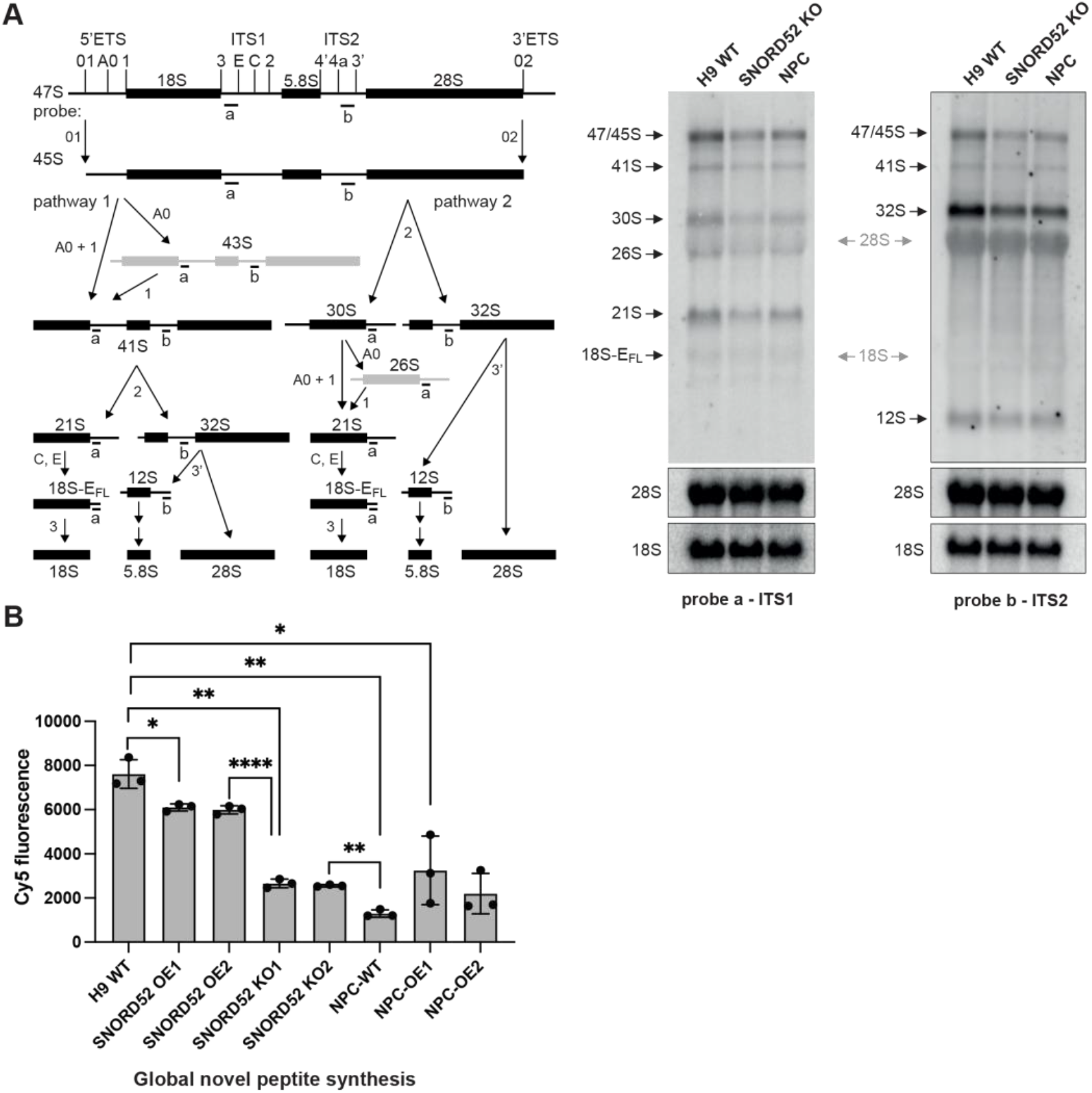
Analysis of ribosome biogenesis, global translation, and polysome profiles in cells lacking 28S:3904-me. A: Analysis of rRNA biogenesis pathways in H9^WT^ (WT), H9^52KO^ (SNORD52 KO) cells, and H9^NPC^ (NPC). Left: schematic showing rRNA processing steps and location of probes ‘a’ and ‘b’. Right: levels of pre-rRNA, processing intermediates, and mature 18S and 28S rRNA as assessed by northern blot. B: Nascent peptide synthesis assessed by OPP incorporation assay in H9^WT^ (WT), H9^WT^ with an EGFP-only transgene (WT-EGFP), H9^52KO^ (KO1 and KO2) cells, H9^NPC^ (NPC-WT), WT-EGFP derived NPCs (NPC-WT-EGFP), and H9^OE-NPC^ cells (NPC-OE1 and NPC-OE2). Equal amounts of cells were seeded. Cy5 fluorescence (Y-axis) is proportional to peptide synthesis. SD values refer to 3 independent experiments. ns: not significant. *: P≤0.05, **: P≤0.01, ***:P≤0.001, ****:P≤0.0001 (Welch’s unpaired t-test). C: Polysome profiles for H9^WT^ (WT, black), H9^52KO^ (52-KO1, red), and H9^NPC^ cells (NPC, blue). Average values from three independent experiments. The Y-axis depicts the absorption at A260, the X-axis the time of flow-through.

**Supplementary Figure 8:**
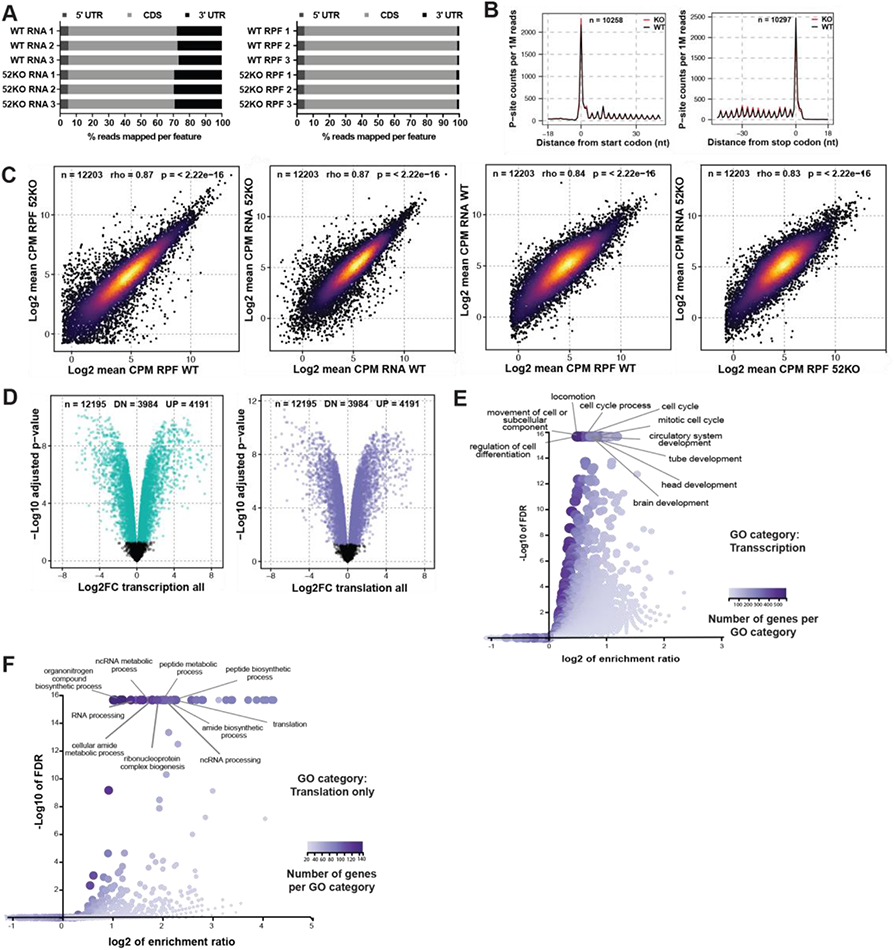
Identification of transcriptional and translational differences between H9^WT^ and ^H952KO^ cells by ribosome profiling. A: Percentage of reads mapping to 5′ untranslated regions (5′ UTR), coding sequences (CDS), or 3′ untranslated regions (3′ UTR) of protein-coding genes, for each replicate sequencing library of H9^WT^ (WT) and H9^52KO^ (52KO) cell lines. Both total RNA (left) and ribosome protected fragment (RPF, right) derived reads are shown. B: Periodicity of ribosome protected fragment (RPF) reads mapped to mRNA transcripts. Metagene analysis shows normalized mean counts, at single-nucleotide resolution, representing ribosome P-site occupancy relative to start (left) and stop (right) codons from H9^WT^ (WT, black) or H9^52KO^ cells (KO, red) libraries (*n* = 3 libraries from individual cultures). Number of transcripts analyzed after extreme outlier removal is given by ‘n’. C: Correlation between reads mapped per mRNA transcript in total RNA and RPF libraries in H9^WT^ (WT) and H9^52KO^ (KO) (*n* = 3 libraries from individual cultures). Normalized (CPM) mean counts are plotted. Color scale indicates plotting density. ‘n’ denotes number of transcripts analyzed. Spearman’s rho and associated *P* value (algorithm AS 89) are shown. D: Left: mRNA transcripts displaying altered expression level (Log2FC mRNA) in H9^52KO^ (KO) compared to H9^WT^ (WT) as measured by analysis of read counts mapped to mRNA transcripts derived from total RNA libraries. Those changing significantly between conditions are colored (cyan). Right: differential ribosome occupancy on mRNA transcripts in H9^52KO^ compared to H9^WT^ cells. Log2 fold-change in read counts derived from analysis of RPF libraries (Log2FC RPF) and corresponding −Log10 of Benjamini-Hochberg *P^adj^* values are plotted. Transcripts changing significantly between conditions are colored (purple). ‘n’ denotes total number of transcripts analyzed. The number of transcripts showing reduced (DN) or increased (UP) translation is also shown. E: Gene ontology analysis of mRNA transcripts displaying upregulated expression in H9^52KO^ cells compared to H9^WT^ cells. Top 10 GO categories (FDR < 0.05) for biological processes are listed. Number of genes overlapping with each biological process GO category is indicated by the color scale gradient (count). F: Gene ontology analysis of mRNA transcripts displaying decreased translation (ribosome occupancy) but not transcription (translation only) in H9^52KO^ cells compared to H9^WT^. Top 10 GO categories (FDR < 0.05) for biological process are labelled. Number of genes overlapping with each biological process GO category is indicated by the color scale gradient (count).

**Supplementary Figure 9:**
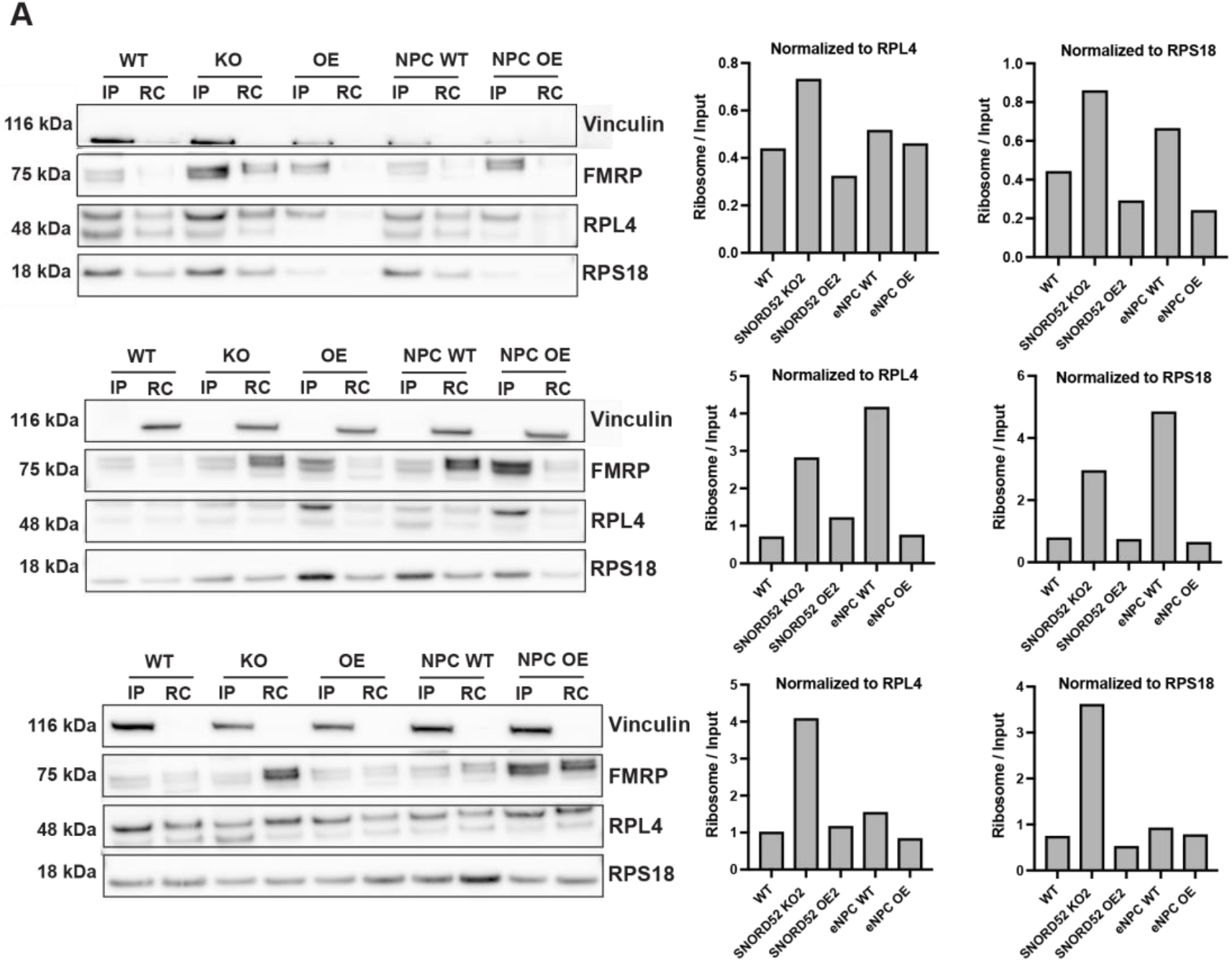
Analysis of the potential molecular mechanisms underlying differential translation by wild-type and SNPRD52 KO cells. A: Complement to Figure 6C and D, 3 more experimental repeats. Left: Detection of total cellular and ribosome-bound FMRP protein by Western Blot in H9^WT^ (WT), H9^52KO^ (KO), H9^52OE^ (OE), H9^NPC^ (NPC WT), and H9^52OE-NPC^ (NPC OE) cells. IP (input) = whole cell lysate, RC (ribosome cushion) = ribosomes purified by sucrose cushion. Vinculin serves as a control for appropriate separation of input and purified ribosomes, RPL4 and RPS18 for normalization to ribosome abundance. Right: Quantification of the enrichment of FMRP in the ribosome-bound fraction based on western blots (left). Bands were quantified, then subsequently the values for FMRP normalized to the corresponding ribosomal protein (RPL4 or RPS18), and finally the ratio of the ribosome-bound fraction over the input fraction calculated.

## SUPPLEMENTARY TABLES

**SUPPLEMENTARY TABLE 1**

rRNA positions displaying statistically significant 2’-O-me changes during mouse cortex development. RMS values for individual libraries (replicates), average RMS values (mean RMS Score), and corresponding standard deviations (SD RMS score) are listed. ΔRMS refers to the difference in RMS score between the maximal (MAX) and minimal (MIN) average RMS values of the series. The p-value was calculated by a two-tailed Welch’s t-test (assuming unequal variance). The cutoff for ΔRMS is >0.15 and the positions are ranked by p-value.

**SUPPLEMENTARY TABLE 2**

rRNA positions with statistically significant differences in RMS score (as defined for Supplementary Table 1) between four brain regions (cortex CTX, olfactory bulb OFB, cerebellum CBM, and hippocampus HPC) in neonates (P0).

**SUPPLEMENTARY TABLE 3**

rRNA positions with statistically significant differences in RMS score (as defined for Supplementary Table 1) between four brain regions (cortex CTX, olfactory bulb OFB, cerebellum CBM, and hippocampus HPC) in adult mice.

**SUPPLEMENTARY TABLE 4**

rRNA positions with statistically significant differences in RMS score (as defined for Supplementary Table 1) between adult mice and neonates (P0) in four brain regions (cortex CTX, olfactory bulb OFB, cerebellum CBM, and hippocampus HPC).

**SUPPLEMENTARY TABLE 5**

rRNA positions with statistically significant differences in RMS score (as defined for Supplementary Table 1) between H9^WT^ hESCs and their differentiated progeny respectively (ectoderm, endoderm, mesoderm).

**SUPPLEMENTARY TABLE 6**

List of rRNA positions considered to be true potentially 2’-O-methylated positions in this article for mouse and human based on RMS and MassSpec evidence. The numbering is based on the snoRNABAse (snorna.biotoul.fr).

**SUPPLEMENTARY TABLE 7**

Ribosome-associated peptides identified by mass spectrometry from H9^WT^ hESCs, H9^52KO^, and H9^eNPC^ cells as well as either 80S monosomes or polysomes and compared two by two.

**SUPPLEMENTARY TABLE 8**

Peptides described in Supplementary Table 8, filtered by a list of true ribosome associated proteins (Supplementary Table 9).

**Supplementary Table 9**

List of true ribosome associated proteins described by Imami et al.

**Supplementary Table 10**

List of canonical transcripts representing each protein-coding gene selected from the GRCh38, v97 Ensembl annotation file.

## Methods

### Cell culture

#### Human embryonic stem cell (hESC) and induced pluripotent stem cell (iPSC) culture

H9 and HUES4 cells were grown under feeder-free conditions on plates coated with hESC-qualified Matrigel (Corning Life Sciences, #354277) in mTeSR™1 medium (Stem Cell Technologies, #85850). Medium changes were performed daily. Cells were passaged about every three days using 1X TrypLE^TM^ Select (Life Technologies, #12563-011) when reaching 80-90% confluence.

Upon thawing, Rock inhibitor (Y-27632) (LC laboratories, #Y-5301) was added at a final concentration of 10µM. Cells were frozen in half mTeSR™1 medium and half FBS-20% DMSO (final DMSO concentration: 10%).

KOLF2 iPS cells were grown under feeder-free conditions on Matrigel-coated plates in TeSR™-E8™ (Stem Cell Technologies, #05990).

#### Generation of neural progenitor cells

H9 hESCs were differentiated into early neural progenitor cells (eNPCs) with STEMdiff^TM^ Neural Induction Medium (Stem Cell Technologies, #05835) following the manufacturer’s instructions for the monolayer protocol variant. In brief, H9 cells were plated on Matrigel-coated plates at a density of 2×10^6^ cells per cm in neural induction medium supplemented with 10µM Rock inhibitor. Medium was changed every day and the cells passaged after about a week. After two more passages in neural induction medium, the cells were transferred into STEMdiff^TM^ Neural Progenitor Medium (Stem Cell Technologies, #05833) for expansion and freezing.

#### Long-term neural differentiation

H9-derived NPCs were further differentiated into late neural progenitor cells (lNPCs, or neural precursors, according to the manufacturer) by plating them at 125.000 cells/cm^2^ on Matrigel and growing the cells in STEMdiff™ Neuron Differentiation Kit (Stem Cell Technologies, #08500) medium for one week with daily medium changes.

After one week, the cells were dissociated and plated at a density of 3×10^4^ cells/cm in STEMdiff™ Neuron Maturation Kit (Stem Cell Technologies, #08510) medium on a double layer of poly-L-ornithine (Sigma-Aldrich #P3655) at 15µg/mL in PBS and laminin (Sigma-Aldrich, #L2020) at 5µg/mL in DMEM/F-12. The medium was changed every second day and the cells kept in culture up to 52 days.

#### Neural progenitor cell maintenance

H9 hESC-derived early neural progenitor cells (eNPCs) were grown on Matrigel (Corning Life Sciences, #354277) in STEMdiff^TM^ Neural Progenitor Medium (Stem Cell Technologies, #05833).

Cells were passaged around every five days at 80-90% confluence using Accutase (Stem Cell Technologies, #07920) for detaching and DMEM/F-12 + GlutMAX (Invitrogen, #31331-028) for resuspension, and frozen in STEMdiff^TM^ Neural Progenitor Freezing Medium (Stem Cell Technologies, #05838).

#### Directed ES differentiation into the three embryonic germ layers

HUES4, H9, and KOLF2 iPS cells were differentiated into ecto-, endo-, and mesoderm using the STEMdiff^TM^ Trilineage Differentiation Kit (Stem Cell Technologies, #05230) according to the manufacturer’s instructions.

In brief, ES cells were plated on day 0 in technical triplicates in Matrigel-coated 6-well plates in either mTeSR™1 (HUES4, H9) or TeSR™-E8™ medium (KOLF2 iPS) supplemented with 10µM Rock inhibitor for endo-and mesoderm differentiation at 2.10^6^ cells per well for endoderm and 0,5.10^6^ cells per well for mesoderm. Cells destined for ectoderm differentiation were directly plated in ectoderm differentiation medium on day 0, supplemented with 10µM Rock inhibitor, at a density of 2.10^6^ cells per well. On day 1, medium was switched to the respective germ layer differentiation medium and changed on a daily basis until day 5 for endo- and mesoderm differentiation, and day 7 for ectoderm differentiation. Bright field images were taken every day and cells from every day of differentiation analyzed by RT-qPCR and immunohistochemistry.

### RNA isolation from mouse brains

CD-1 mice were sacrificed at embryonic stages E11, E12.5, E14, E15.5, E17, P0 (neonates) and 6-7 months (adult). The brains were dissected in ice cold PBS in a cold room and snap-frozen on dry ice. RNA was extracted using TRIZOL-LS according to the manufacturer’s instructions.

### RiboMeth-Seq

RiboMeth-seq library construction and sequencing were performed as previously described (Birkedal et al., 2014; Krogh et al., 2016). Triplicate libraries were produced for each cell line or condition analyzed, and grown to ∼70–80% confluence before collection. A portion of 5 μg of total RNA was used for input. RNA was partially degraded in alkali at denaturing temperatures. The 20–40-nucleotide fragments were purified by PAGE and linkers added using a system relying on a modified *Arabidopsis* tRNA ligase joining 2′,3′-cyclic phosphate and 5′-phosphate ends. The libraries were sequenced on the Ion Proton platform using Ion PI Chip Kit v.3 (Life Technologies).

### RiboMeth-seq Data treatment

Data was analyzed as previously reported (Birkedal et al., 2014; Krogh et al., 2016). Briefly, sequencing reads were mapped to a corrected human rRNA reference sequence. To facilitate comparison with other studies, we have used the human rRNA sequence numbering according to snoRNABase throughout this study (snorna.biotoul.fr). An alignment table of these rRNA sequences is provided in Krogh and Jansson et al, 2016 (Krogh et al., 2016). The RiboMeth-seq score (RMS score) represents the fraction of molecules methylated at each nucleotide position, and is calculated by comparing the number of read-end counts at the queried position to six flanking positions on either side. Quantifications are performed in mouse on 41 sites in 18S, 66 in 28S and 2 in 5.8S and in human on 41 sites in 18S, 68 in 28S and 2 in 5.8S respectively, for which both RMS plus mass spectrometry evidence exists, and are reliably detected in at least one of cell lines examined in this study (Supp. Table 6). As to facilitate comparison, the human numbering is used both for mouse and human samples. The equivalence between mouse and human sites can be found in Supplementary Table 6.

Statistically significant differences in RMS signatures between two cell lines or conditions were determined by pairwise comparison (p<0.05, two-tailed unpaired Welch’s t test and ≥0.15 difference in RMS score).

Heat-map representations were produced using the pheatmap function in R.

RMS data has been deposited to Gene Expression Omnibus (GEO), accession: **GSE205022**

### Generation of loss and gain of function mutants

#### Generation of SNORD52 knock-out clones by CRISPR-Cas9

Two guides (sgRNAs) encompassing the human SNORD52 gene were designed: GGAGTGGACGTTAGAAAGGG and GGATACTTGGGTCTCCAGAA.

Both sgRNAS were cloned into pX335 and pX458 plasmids respectively. The plasmids were co-transfected into H9 hESC cells using the Amaxa 4D nucleofector (#AAF-1003B and #AAF-1003X) and the P3 Primary Cell 4D-Nucleofector X kit (Lonza, #V4XP-3024).

48h after transfection, the cells were single-cell sorted into Matrigel-coated 96-well plates for double GFP/Crimson fluorescence by FACS. Rock inhibitor (Y-27632) was added to mTESR-1 medium until the second medium change.

After about two weeks, colonies became visible and were screened for successful deletion using standard PCR (forward primer: CTCCCAGTGGAGCTGTTCTC, reverse primer: GGGGGAGATTCCAAACCTTA), the GoTaq Green master mix (Promega, #M7122), and running the amplification products on a 1% Agarose gel.

Candidate clones were further verified by DNA sequencing and expanded. A few CRISPR-negative clones – showing no deletion of SNORD52 but having undergone otherwise the exact same procedure – were also expanded and tested alongside the two SNORD52 KO clones.

#### Generation of SNORD52 overexpression clones

The SNORD52 gene sequence was cloned into an artificial intron and placed into a 2-exon EGFP sequence derived from pGINT (courtesy to Cristian Bellodi), subsequently used to replace the intron-less EGFP of the AAVS1-targeting vector pAAV-PuroCAG-EGFP obtained from Ludovic Vallier.

Following the protocol described in Bertero et al. (Bertero et al., 2016), the EGFP-SNORD52 construct was targeted to the AAVS1 safe harbor locus in H9 hESCs *via* zinc finger nucleases. As a control, the original pAAV-PuroCAG-EGF construct was used.

H9 hESCs were transfected with the Amaxa 4D Nucleofector. The cells were expanded for a few days, allowing for the elimination of transient transfection, then single cell FACS-sorted for GFP fluorescence. Correct insertion of the construct into the AAVS1 locus was verified by DNA sequencing, and SNORD52 expression by RT-qPCR.

### Gene expression

#### RNA isolation

Total RNA preparation was performed using QIAZOL (Qiagen) and chloroform according to the protocol from the manufacturer. Concentrations were measured using a NanoDrop, and RNA quality assessed with a BioAnalyser.

#### cDNA generation and RT-qPCR

Reverse-transcription to cDNA was achieved with the TaqManReverese Transcription Kit (Applied Biosystems, #N80803234) according to the manufacturer’s recommendations. Typically, 1µg of RNA was used per reverse transcription reaction.

RT-qPCR analyses were performed in technical triplicates on a StepOnePlus^TM^ Real-Time PCR System (Thermo Fisher Scientific, #4376600) in a 96-well format and using the Fast SYBR Green Master Mix (Thermo Fisher Scientific, #4385612). 2µL of cDNA were combined with 5µL of Fast SYBR Green Master Mix, 0.3µL of forward or reverse primer at 100µM, and 2.4µL nuclease-free water per well.

List of primers used in the study:

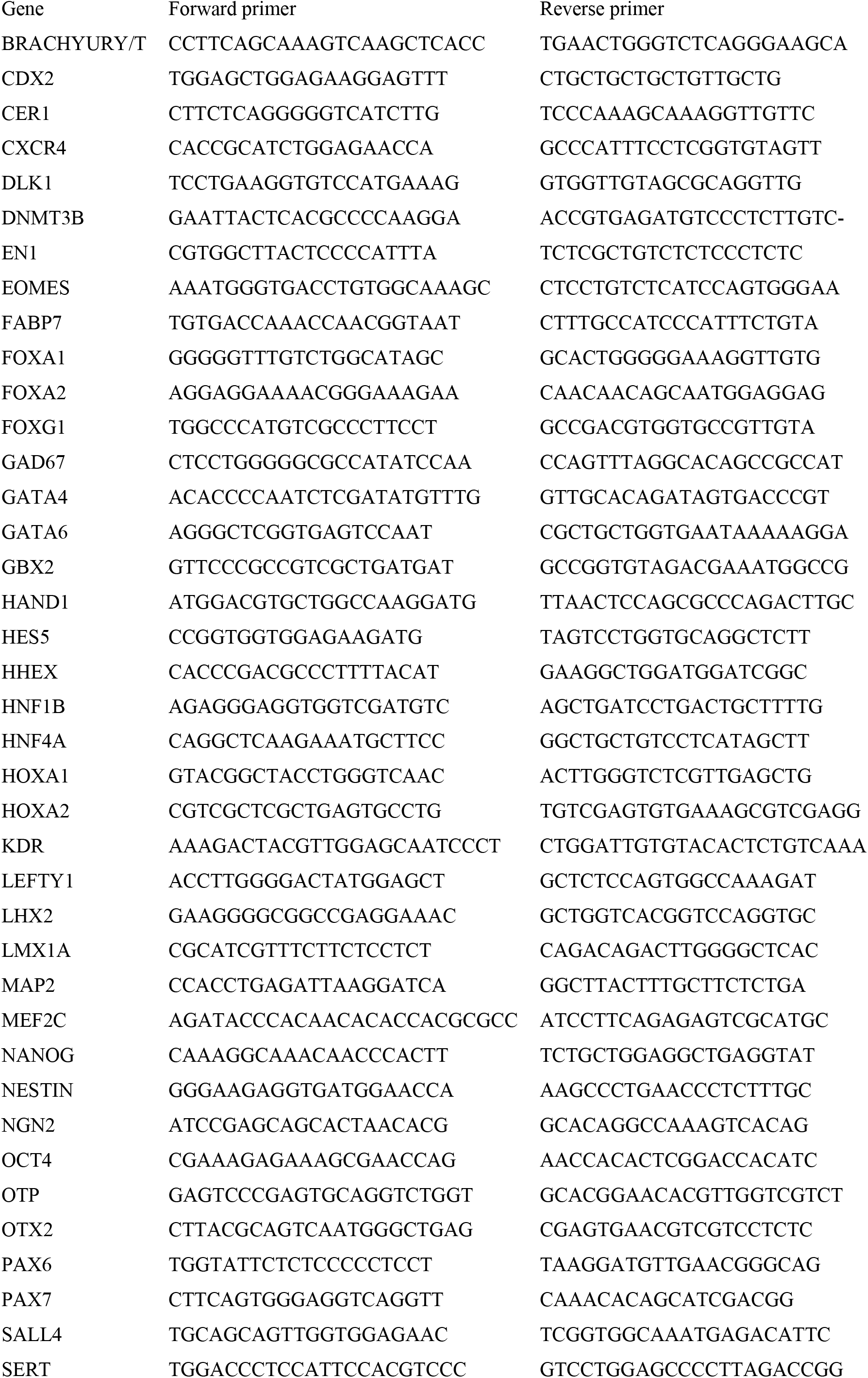
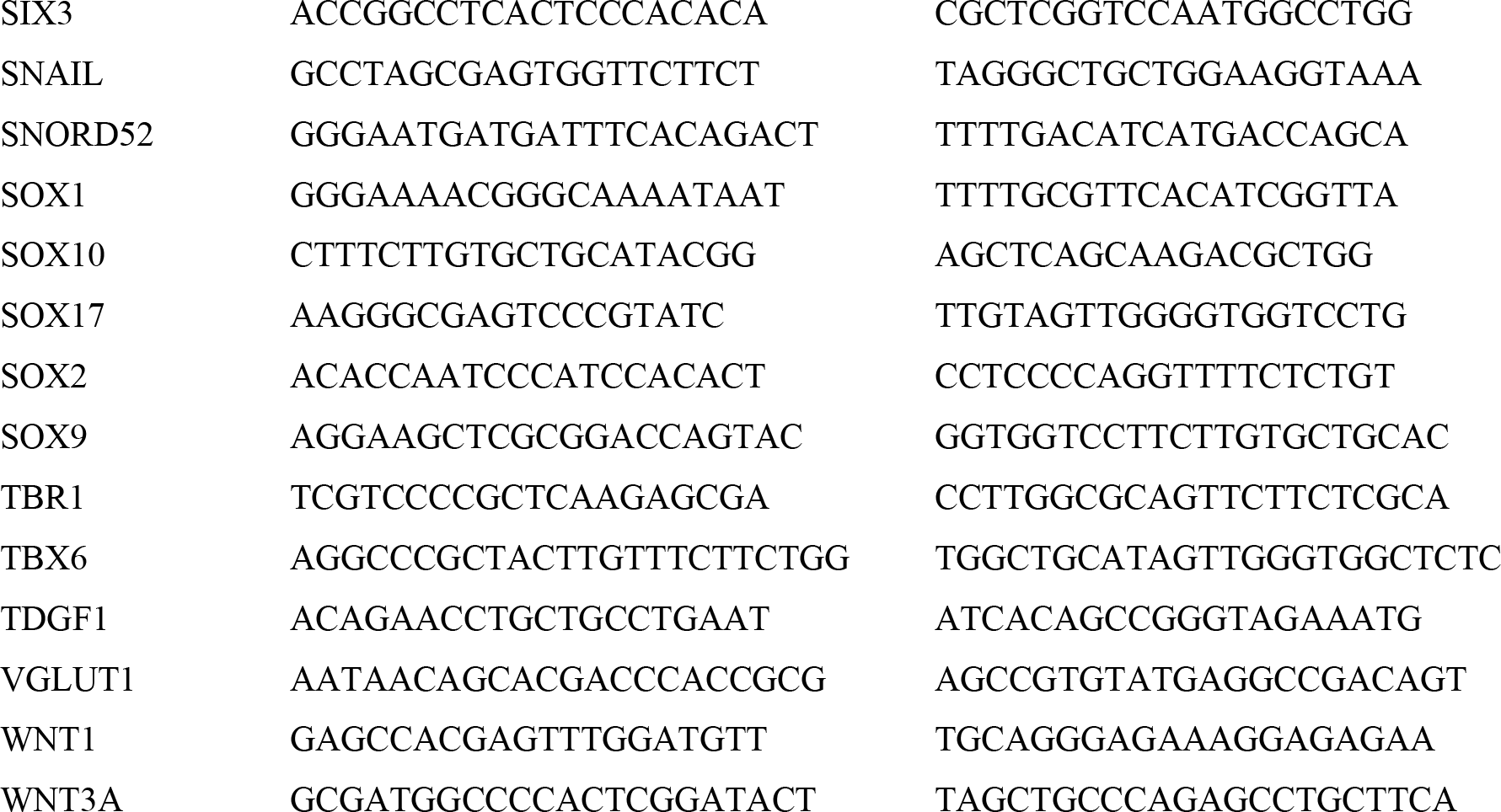

#### Immunofluorescence

Cells were plated in Lab-Tek Chamber slides (Sigma Aldrich, #C7182) at the densities corresponding to the relevant protocol, washed with PBS and fixed for 15 minutes at RT with 4% paraformaldehyde, followed by three PBS washes and potential storage in PBS at 4°C. Fixed cells were permeabilized for 10 minutes in PBS, 0.1% Triton, then incubated for 1h in blocking buffer (PBS, 5% FBS). Primary antibodies were diluted in blocking buffer at the indicated dilutions and left overnight at 4°C under gentle shaking, then washed off by applying PBS 3 times for 10 minutes. Secondary antibodies were diluted at a 1:1000 dilution in blocking buffer and left for 1h at room temperature. Slides were subsequently washed 3 times with PBS for 10 minutes and mounted in Duolink in situ mounting medium with DAPI (Sigma-Aldrich, #DUO82020-5ML).

Images were acquired with a Zeiss Axio Imager.M2 microscope (#490020-0004-000) and images analyzed with the open-source ImageJ software (Fiji).

List of primary antibodies:

| Target | Provider | Reference | Host | Isotype | Working dilution |
| --- | --- | --- | --- | --- | --- |
| OCT4 | Stem Cell Technologies | #60093 | Mouse | IgG2b, kappa | 1:1000 |
| PAX6 | Stem Cell Technologies | #60094 | Rabbit | IgG, polyclonal | 1:500 |
| ZO1 | BD Biosciences | #610966 | Mouse | IgG1 | 1:500 |
| TUJ1 | Stem Cell Technologies | #60092 | Mouse | IgG2a | 1:500 |
| MAP2 | Abcam | #ab112667 | Mouse | IgG1 | 1:500 |
| NESTIN | Stem Cell Technologies | #60091 | Mouse | IgG1, kappa | 1:2000 |
| OTX2 | Abcam | #ab21900 | Rabbit | IgG, polyclonal | 1:500 |
| TTR | BioRad | #AHP1837 | Sheep | IgG, polyclonal | 1:500 |
| SOX17 | R&D Systems | #AF1924 | Goat | IgG, polyclonal | 1:400 |
| T/BRACHYURY | R&D Systems | #AF2085 | Goat | IgG, polyclonal | 1:200 |
| Beta-catenin | Cell Signaling | #9587 | Rabbit | IgG, polyclonal | 1:500 |

List of secondary antibodies:

| Antibody | Provider |
| --- | --- |
| Goat anti-Mouse IgG (H+L) Highly Cross-Adsorbed Secondary Antibody, Alexa Fluor 488 | Thermo Scientific |
| Goat anti-Mouse IgG (H+L) Highly Cross-Adsorbed Secondary Antibody, Alexa Fluor 594 | Thermo Scientific |
| Donkey anti-Goat IgG (H+L) Cross-Adsorbed Secondary Antibody, Alexa Fluor™ 594 | Thermo Scientific |
| Donkey Anti-Sheep IgG H&L (Alexa Fluor® 594) | Abcam |
| Goat anti-Rabbit IgG (H+L) Highly Cross-Adsorbed Secondary Antibody, Alexa Fluor 488 | Thermo Scientific |
| Donkey anti-Rabbit IgG (H+L) Highly Cross-Adsorbed Secondary Antibody, Alexa Fluor 594 | Thermo Scientific |

### Western Blotting

#### Normal Western Blot

Western blotting was performed as previously described (Jansson et al., 2015). The following antibodies were used:

| Target | Provider | Reference | Host | Isotype | Working dilution |
| --- | --- | --- | --- | --- | --- |
| RECK | Cell Signaling | #3433 | Rabbit | Monoclonal | 1:1000 |
| CITED1 | Merck | #424M-17 | Mouse | Monoclonal | 1:500 |
| Vinculin | Sigma Aldrich | #V9131 | Mouse | Monoclonal | 1: 100 000 |
| FMRP | Cell Signaling | #4317 | Rabbit | Polyclonal | 1:1000 |
| RPL4 | Santa Cruz Biotechnology | #sc-100838 | Mouse | Monoclonal | 1:1000 |
| RPS18 | Nordic Biosite | #LS-C162774-400 | Rabbit | Polyclonal | 1:1000 |

#### Ribosome isolation by sucrose cushion followed by Western Blot

### 3D modelling

Modelling performed using MacPyMol (Version 2.1 INTEL-12.10.12) on the structure 4UG0 (https://www.rcsb.org/structure/4UG0) published by Khatter et al. (Khatter et al., 2015).

for the FMRP part:

Structure published by Chen et al. (Chen et al., 2014).

Accession number: EMD-5806 on http://emsearch.rutgers.edu.

### Ribosome profiling

#### Ribosome profiling

Ribosome Profiling was performed essentially as previously described (Ingolia et al., 2009), following the protocol given in TruSeq Ribo Profile Mammalian (Illumina), with minor modifications. Three individual replicates for each of the two cell lines were collected. A single 15cm dish corresponding to one replicate was harvested at a time. For each replicate, cell media was aspirated and cells washed with ice-cold PBS. No cycloheximide pre-treatment was performed. After thorough removal of the PBS, the dish was fully immersed in liquid nitrogen and placed on dry ice. For cell lysis, 1mL of 1x Mammalian Lysis Buffer (Illumina) containing 100 μg/mL cycloheximide was added dropwise to the dish which was then placed on wet ice. Cells were then scraped off to the lower portion of the dish and allowed to thaw in the lysis buffer. Lysate was homogenized by pipetting and triturated ten times through a 25-gauge needle. The lysate was then transferred to a DNA LoBind 1.5mL microfuge tube (Eppendorf) and incubated on ice for 5min. The lysate was cleared by centrifugation at 20000g, 4°C for 10min and the supernatant transferred to a fresh microfuge tube. Aliquots were prepared for each replicate, flash-frozen in liquid nitrogen and stored at −80°C until further use. The steps detailed in TruSeq Ribo Profile Mammalian protocol (Illumina) were followed to generate total RNA and ribosome protected fragment (RPF) RNAseq libraries corresponding to the 3 individual replicates from each of the two cell lines. For RPFlibraries, following nuclease digestion, monosomes were purified using illustra MicroSpin S-400 HR Columns. Ribo-Zero™ Gold Kit (Illumina) was used to deplete ribosomal rRNA. The libraries prepared from total RNA or RPF for both conditions were pooled and sequenced on a NextSeq® 500 System (Illumina).

#### Ribosome Profiling data analysis

The sequencing data was demultiplexed using Illumina bcl2fastq. Quality of the sequencing files were controlled with fastqc. Adaptor sequences were removed with cutadapt. Reads derived from RPF and total RNA were aligned to human rRNA and tRNA sequences with bowtie2 (v2.2.9) and the mapped reads discarded. The remaining reads were aligned to GRCh38.p12 (Ensembl v.97) with STAR. Reads mapping to Human Genome Organisation (HUGO) approved genes were used for downstream analyses. RPF read lengths were analysed for trinucleotide periodicity using Ribotaper. RPF reads with lengths between 29 to 34 nucleotides were selected and the optimal P-site offset was defined as position 12 from 5’ read ends. RPF read alignment files were filtered with samtools to retain only 29 to 31nt read lengths, no read length filtering was applied to the total RNA alignment files. A single canonical transcript representing each protein-coding gene was selected from the GRCh38, v97 Ensembl annotation file (Supp. Table 10). 649 FeatureCounts was used to generate counts of primary reads mapping to exons of these transcripts for both total RNA and RPF. RPF reads with ribosome P-site positions mapping within transcript coding-region sequences (CDS) were again counted using FeatureCounts, and along with the mRNA exons mapped reads, used for further measurements of differential translation and mRNA expression. Ribosomal Investigation and Visualization to Evaluate Translation (RIVET) 49, was used for translation and expression analysis of the representative transcripts (similar results were obtained for gene-level analysis). No fold-change cut-offs were directly applied, in order to additionally detect more subtle changes in translation. Regulated transcripts were therefore nominally identified by statistical significance. Translation regulation categories were defined according to RIVET, based on mRNA expression and ribosome occupancy, derived from normalized total RNA read counts or RPF read counts mapping to protein-coding mRNA transcripts, respectively (Supplementary Table 2). Plots from the resulting RIVET output files were generated using the ggplot2 package in R. The RNA sequencing data has been deposited to GEO, accession: **GSE199387.**

#### Gene ontology and gene-set enrichment analysis

All Gene ontology (GO) analyses of ribosome profiling was performed using the WebGestalt using the over-representation test against GO biological process database (Liao et al., 2019).

#### Metagene analysis

For metagene analyses, bam files containing exon-mapped reads for each library were converted to normalized reads per kilobase per million (RPKM) or counts per million reads (CPM) single-nucleotide resolution coverage bigwig files, with bamCoverage from the deepTools suite60. WiggleTools61 (Ensembl) and wigToBigWig (Encode, kentUtils) were then used to merge these and create mean coverage files per condition. These were input to deepTools computeMatrix, together with an annotation file containing the exon coordinates for the selected mRNA transcripts. For RPF coverage over all transcripts, a count matrix was then generated for library RPKM RPF coverage over the coding regions (CDS), scaled to size 100 nt, flanked by unscaled regions before and after the translation start (TSS) and end (TES) sites. For further analysis, the scaled coverages of transcripts comprising the different translationally regulated categories were extracted from this matrix and median values at each position plotted. For average ribosome occupancy, CPM normalization was used and offset applied using bamCoverage, so as to use only the nucleotide position representing the ribosome P-site for each read as the signal (see ‘Ribosome profiling data analysis’, above). The P-site coverage files were input to computeMatrix and a count matrix generated for −30 to +330 or −330 to +30 nucleotides, relative to the CDS start or end site respectively for each transcript (unscaled). The resulting counts at each position were divided by the total RPF count in CDS for each corresponding transcript to give the average ribosome occupancy per nucleotide position in each transcript. The mean values at each equivalent nucleotide position relative to the translation start site were plotted after extreme outlier removal (>3x interquartile range), no smoothing was applied. For P-site CPM the same matrices were used, although here the counts at each position were summed at each nucleotide position. For plotting, extreme outliers (>3x interquartile range) were removed. Plots were produced using ggplot2 in R.

### Polysome profiling

Cells at 70-80% confluency were incubated with 100 µg/mL cycloheximide (Sigma Aldrich) for 3 min., washed once, and harvested by scraping in PBS containing 100 µg/mL cycloheximide. Cells were lysed at 4°C for 10 min in polysome buffer (20 mM Tris-HCl (pH 7.5), 150 mM KCl, 5 mM MgCl_2_) supplemented with 0.5% NP40 (Igepal CA-630, Sigma Aldrich), 2 mM DTT, 100 µg/mL cycloheximide, Protease Inhibitor Cocktail (cOmplete EDTA free, Roche), and murine RNase inhibitor (NEB). The cell lysate was cleared for membranes by centrifugation at 12.000g for 15 min. at 4°C. Cleared lysates were normalized according to NanoDrop UV spectrophotometer measurements (Thermo Scientific) and layered onto a 7% −47% (w/v) linear sucrose gradient (Sigma BioUltra) in polysome buffer. Gradients were centrifuged at 35.000 g for 3h at 4°C in an ultracentrifuge using the SW40ti rotor head (Beckman). Fractions of 1 mL were collected from the top at 1 mL/min. while continuously measuring A_254_ using a Brandel Tube Piercer (Brandel) and the BioLogic LP system (BioRad).

### Global nascent peptide synthesis assay

Global novel protein synthesis was assessed using the Click-iT Plus OPP Protein Synthesis Assay Kit (Life Technologies, #C10456 for green fluorescence, #C10457 for red fluorescence given that the overexpression clones are constitutively expressing EGFP) according to the manufacturer’s protocol.

### TOP/FOP Wnt reporter assay

The M50 Super 8x TOPFlash (Addgene #12456) and M51 Super 8x FOPFlash (Addgene, #12457) were transfected together with a pRLTK (Renilla) plasmid using the Mirus2020 transfection reagent (MirusBio, #MIR 5404) according to the supplier’s protocol in a 12-well plate format in technical triplicates.

Cells were prepared for Luciferase and Renilla measurements using the Dual-Glo Luciferase Assay System (Promega, #E2920) according to the manufacturer’s instructions. Luminescence readings were carried out on a GloMax Multi Detection System (Promega). Luciferase readings were normalized to the corresponding Renilla values.

### Northern blotting analysis of rRNA processing

Total RNA (7.5 μg) was separated on a formaldehyde denaturing 1% agarose gel and transferred to a BrightStar-Plus membrane (Ambion) using capillary blotting. followed by UV cross-linking. The probes (10 pmol each) were radiolabelled with [γ-32P]ATP using T4 PNK (Thermo Scientific) and hybridized to the membrane overnight in hybridization buffer (4× Denhardt’s solution, 6× SSC, 0.1% SDS) at *T*m of the probe of −10 °C. The membrane was subsequently washed four times in 3× SSC supplemented with 0.1% SDS, followed by exposure to a propidium iodide screen and scanned on a Typhoon scanner (GE Healthcare).

### Mass spectrometry on 80S and polysomes

Proteins from polysome profile fractions containing 80S ribosomes and polysomes respectively, were precipitated with 20% Trichloroacetic Acid and washed three times in ice cold acetone. Protein pellets were solubilized using 100 µl of lysis buffer (50 mM HEPES (pH 8.5), 6 M guanidinium hydrochloride, 10 mM TCEP, 40 mM CAA). Samples were boiled at 95°C for 5 min., after which they were sonicated on high for 5×30 sec. in a Bioruptor sonication water bath (Diagenode) at 4°C. After determining protein concentration with Bradford (Sigma), 10 µg was taken forward for digestion. Samples were diluted 1:3 with 10% Acetonitrile, 50 mM HEPES pH 8.5, LysC (MS grade, Wako) was added in a 1:50 (enzyme to protein) ratio, and samples were incubated at 37°C for 4 hrs. Samples were further diluted to 1:10 with 10% Acetonitrile, 50 mM HEPES (pH 8.5), trypsin (MS grade, Promega) was added in a 1:100 (enzyme to protein) ratio and samples were incubated overnight at 37°C. Enzyme activity was quenched by adding 2% trifluoroacetic acid (TFA) to a final concentration of 1%. Prior to TMTPro labeling, the peptides were desalted on in-house packed C18 Stagetips (Rappsilber et al., 2007). For each sample, 2 discs of C18 material (3M Empore) were packed in a 200ul tip, and the C18 material activated with 40 µl of 100% Methanol (HPLC grade, Sigma), then 40 µl of 80% Acetonitrile, 0.1% formic acid. The tips were subsequently equilibrated 2× with 40 µl of 1% TFA, 3% Acetonitrile, after which 10 µg of sample was loaded using centrifugation at 4,000 rpm. After washing the tips twice with 100 µl of 0.1% formic acid, the peptides were eluted into clean 500 µl Eppendorf tubes using 40% Acetonitrile, 0.1% formic acid. The eluted peptides were concentrated in an Eppendorf Speedvac, and re-constituted in 50 mM HEPES (pH 8.5) for TMTPro labeling. For normalization, an equimolar peptide mix from all the samples was generated by mixing equal amounts from each sample, that could subsequently act as a normalization spike-in. Labeling was done according to manufacturer’s instructions, and subsequently, labeled peptides were mixed 1:1:1:1:1:1:1:1:1:1:1:1:1 (13-plex, 12 samples + 1 normalization spike-in), acidified to 1% TFA and Acetonitrile concentration brought down to <5% using 2% TFA. Prior to mass spectrometry analysis, the peptides were fractionated using an offline ThermoFisher Ultimate3000 liquid chromatography system using high pH fractionation (5mM Ammonium Bicarbonate, pH 10) at 5 µl/min flowrate. 10 µg of peptides were separated over a 70 min. gradient (5% to 35% Acetonitrile), while collecting fractions in 204 sec. intervals. The resulting 20 fractions were pooled into 10 final fractions and vacuum concentrated to dryness. Fractions were resuspended in 1% TFA, 2% Acetonitrile for MS analysis.

### MS data acquisition

For each fraction, peptides were loaded onto a C18 trap cartridge (ThermoFisher 160454), connected in-line to a 50 cm C18 reverse-phase analytical column (Thermo EasySpray ES803) using 100% Buffer A (0.1% Formic acid in water) at 5 µl/min., using the Ultimate3000 HPLC system. After trap loading, the sample loop was switched out of the flowpath, and peptides were eluted over a 90 min. method ranging from 8% to 60% of Buffer B (80% acetonitrile, 0.1% formic acid) at 200 nl/min.. The Orbitrap Fusion instrument (Thermo Fisher Scientific) was run in an SPS MS3 top speed method with FAIMSPro ion mobility enabled (2 CVs, −50V and −70V). Full MS spectra were collected at a resolution of 120,000, with an AGC target of 100% or maximum injection time of 50 ms and a scan range of 400–1600 m/z. The MS2 spectra were obtained in the ion trap operating at turbo speed, with an AGC target value of 1×10^4^ or maximum injection time of 35 ms, a normalised CID collision energy of 35 and an intensity threshold of 5e3. Dynamic exclusion was set to 60 s, and ions with a charge state <2, >6 or unknown were excluded. From the resulting MS2 scan, 10 precursors were selected for SPS-MS3 analysis, fragmented with a normalised HCD collision energy of 55, and ions collected for a maximum of 118 ms or AGC target of 250%. Resulting MS3 spectra were collected at 60,000 resolution and scan range of 100-500 for reporter ion quantification. FAIMS CVs were switched on the fly, with 1.5 s cycle time dedicated to each CV. MS performance was verified for consistency by running complex cell lysate quality control standards, and chromatography was monitored to check for reproducibility.

The mass spectrometry data have been deposited to the ProteomeXchange Consortium (http://proteomecentral.proteomexchange.org) via the PRIDE partner repository with the dataset identifier **PXD035621**.

### TMT Quantitative Proteomics Analysis

The raw files were analyzed using Proteome Discoverer 2.4. TMT SPS-MS3 quantitation was enabled in the processing and consensus steps, and spectra were matched against the 9606 Human database obtained from UniProt. Dynamic modifications were set as Oxidation (M), Deamidation (N,Q) and Acetyl on protein N-termini. Cysteine carbamidomethyl and TMTPro were set as static modifications on peptide N-termini and Lysine residues. All results were filtered to a 1% FDR, and protein quantitation done using the built-in Minora Feature Detector.

### Proteomic mass spectrometry data analysis

All MS spectra were searched in Proteome Discoverer 2.4 (ThermoFisher), using the SEQUEST algorithm against the human proteome Uniprot database (containing its reversed complement and known contaminants). Spectral matches were filtered to false discovery rate (FDR) <0.01, using the target-decoy strategy combined with linear discriminant analysis. Proteins were quantified only from peptides with an Average Reporter S/N Threshold of 10, and co-isolation specificity of 0.75.

To examine only those proteins most likely to be truly ribosome associated, thus eliminating proteins likely to be contaminants throughout the sucrose gradients, identified proteins were filtered against a complied list of previously identified high confidence 40S/60S interacting proteins *REF1* (Supp. Table 9).

Statistical analyses of protein abundance changes were performed using the DEqMS pipeline for TMT labelled MS data (https://github.com/yafeng/DEqMS) *REF2*, with q.value<0.05. Log2FC changes in protein abundance were determined to be significant at sca.adj.pval (Benjamini-Hochberg method adjusted DEqMS p-values) <0.05 (Supp. Table 7, Supp. Table 8). The abundance values scaled to the reference channel (pooled sample) output from Proteome Discoverer were used as input.

Volcano plots for each comparison were generated using the ggplot2 package in R. Bar charts were generated in Graphpad Prism using DEqMS normalized abundances and statistical significance was tested using two-tailed unpaired Welch’s *t* test.

The mass spectrometry data have been deposited to the ProteomeXchange Consortium, via the PRIDE repository with the dataset identifier **PXD035621.**

## Notes

### Competing Interest Statement

The authors have declared no competing interest.

